# Reduction of insulin/IGF-1 receptor rejuvenates immunity via positive feedback circuit

**DOI:** 10.1101/795781

**Authors:** Yujin Lee, Dae-Eun Jeong, Wooseon Hwang, Seokjin Ham, Hae-Eun H. Park, Sujeong Kwon, Yoonji Jung, Jasmine M. Ashraf, Coleen T. Murphy, Seung-Jae V. Lee

## Abstract

Immunosenescence is considered an inevitable decline in immune function during aging. Here we show that genetic inhibition of the DAF-2/insulin/IGF-1 receptor drastically delays immunosenescence and rejuvenates immunity in *C. elegans*. We find that p38 mitogen-activated protein kinase 1 (PMK-1), a key determinant of immunosenescence, is dispensable for this rejuvenated immunity. Instead, we demonstrate that longevity-promoting DAF-16/FOXO and heat-shock transcription factor 1 (HSF-1) increase immunocompetence in old *daf-2(-)* animals. The upregulation of DAF-16/FOXO and HSF-1 decreases the expression of the *zip*-*10*/bZIP transcription factor, which in turn downregulates INS-7, an agonistic insulin-like peptide, resulting in further reduction of insulin/IGF-1 signaling (IIS). Thus, reduced IIS bypasses immunosenescence and rejuvenates immunity *via* the upregulation of anti-aging transcription factors that modulate an endocrine insulin-like peptide through a positive feedback mechanism. Because many functions of IIS are conserved across phyla, our study may lead to the development of strategies for human immune rejuvenation.

## Introduction

An increase in mortality after infection is a key feature of aging. This age-dependent decline in immunity, i.e., immunosenescence, is nearly universal across phyla (Fulop et al., 2017; Goronzy and Weyand, 2013; Müller et al., 2013). Vertebrates display immunosenescence in adaptive immunity. In addition, old mice and humans have increased pro-inflammatory cytokine levels, termed “inflamm-aging”, because of the chronic activation of innate immune systems (Baylis et al., 2013; Franceschi and Campisi, 2014). However, the complexity of the immune systems and cell types in vertebrates impedes the dissection of the molecular mechanisms underlying immunosenescence.

*Caenorhabditis elegans* is an excellent model for research on aging and immunity, phenomena which are closely related with each other. Many long-lived *C. elegans* mutants display enhanced innate immunity at young age (Alper et al., 2010; Garsin et al., 2003; Yunger et al., 2017; Tiku et al., 2018). Aged *C. elegans* also display an increased susceptibility to pathogen infection, indicating that *C. elegans* can be used as a model of immunosenescence (Kurz and Tan, 2004; Laws et al., 2004). An age-dependent decline in PMK-1/p38 mitogen-activated protein kinase (MAPK) levels underlies *C. elegans* immunosenescence against the pathogenic bacteria *Pseudomonas aeruginosa* (PA14) (Youngman et al., 2011). Although these findings suggest the existence of a link between aging and immunity, the molecular mechanisms by which longevity affects immunosenescence remain largely unknown.

Insulin/insulin-like growth factor 1 (IGF-1) signaling (IIS) plays important roles in aging and immunity in various species, including *C. elegans*. In *C. elegans*, inhibition of IIS, for example, by mutations in *daf*-*2*/insulin/IGF-1 receptor or its agonist, *ins*-*7*, increases the lifespan and enhances resistance against pathogen-mediated death (Evans et al., 2008a; Evans et al., 2008b; Garsin et al., 2003; Kawli and Tan, 2008; Kenyon et al., 1993; Murphy et al., 2003). Reduced IIS activates several transcription factors, such as DAF-16/FOXO, heat-shock transcription factor 1 (HSF-1), and SKN-1/NRF, which upregulate the expression of various genes that contribute to longevity, stress resistance, and immunity (reviewed in Kenyon, 2010; Murphy and Hu, 2013; Riera et al., 2016). Despite extensive research on the roles of IIS in aging and immunity, its role in the regulation of immunosenescence remains unknown.

In this study, we investigated whether longevity interventions affected the rate of immunosenescence in *C. elegans*. Among the various longevity regimens tested here, genetic inhibition of *daf*-*2*/insulin/IGF-1 receptor substantially improved immunocompetence in old worms. We found that temporal inhibition of *daf-2* starting in the middle age rejuvenated immunity in old adult animals. We then showed that delayed immunosenescence conferred by *daf-2* mutations bypassed the functional requirement of PMK-1/p38 MAPK signaling, which is crucial for immunity. Instead, DAF-16/FOXO and HSF-1 increased immunocompetence in old *daf-2* mutants by regulating the expression of various genes. In particular, DAF-16/FOXO and HSF-1 downregulated an agonistic insulin-like peptide, INS-7, *via* decreasing the expression of a bZIP transcription factor, ZIP-10, which in turn reduced IIS further and enhanced immunity at old age. Thus, *daf-2* mutants appear to elude normal immunosenescence *via* a positive-feedback endocrine circuit that consists of the DAF-16/FOXO, HSF-1, and ZIP-10 transcription factors, together with INS-7, which is an agonist of the DAF-2/insulin/IGF-1 receptor.

## Results

### Genetic inhibition of the DAF-2/insulin/IGF-1 receptor improves immunity in aged worms

We wondered whether the rates of immunosenescence were proportionally affected by genetic mutations that promote longevity. Thus, we measured immunosenescence in various long-lived animals, including IIS-defective *daf-2*, sensory-defective *osm-5*, germline-deficient *glp-1*, dietary restriction mimetic *eat-2*, and mitochondrial respiration-impaired *isp-1* mutants. Similar to the wild-type counterparts (Laws et al., 2004; Youngman et al., 2011), *osm-5, glp-1*, and *eat-2* mutants exhibited an age-dependent increase in susceptibility to *Pseudomonas aeruginosa* (PA14) infection (Figures 1A-1E). Old *isp-1* mutants displayed susceptibility to PA14 infection that was comparable to that of young animals; however, their overall survival rate was lower than that of wild-type worms (Figures 1A and 1F). Surprisingly, old (day 9) *daf-2* mutants displayed substantially enhanced survival upon PA14 infection compared with young (day 1) *daf-2* mutants and wild-type worms (Figures 1A and 1G). Consistent with *daf-2* mutant data, old (day 9) *daf-2(RNAi)* worms also displayed enhanced PA14 resistance (Figure 1H), without defects in feeding rates (Figures 1I). Old (day 9) *daf-2* mutant worms also survived longer than did young worms after infection with PAO1, another *P. aeruginosa* strain (Figures 1J and 1K). These data suggest that genetic inhibition of DAF-2 confers sustained immunocompetence against *P. aeruginosa* in old age, which was different from other longevity interventions.

**Figure 1.**
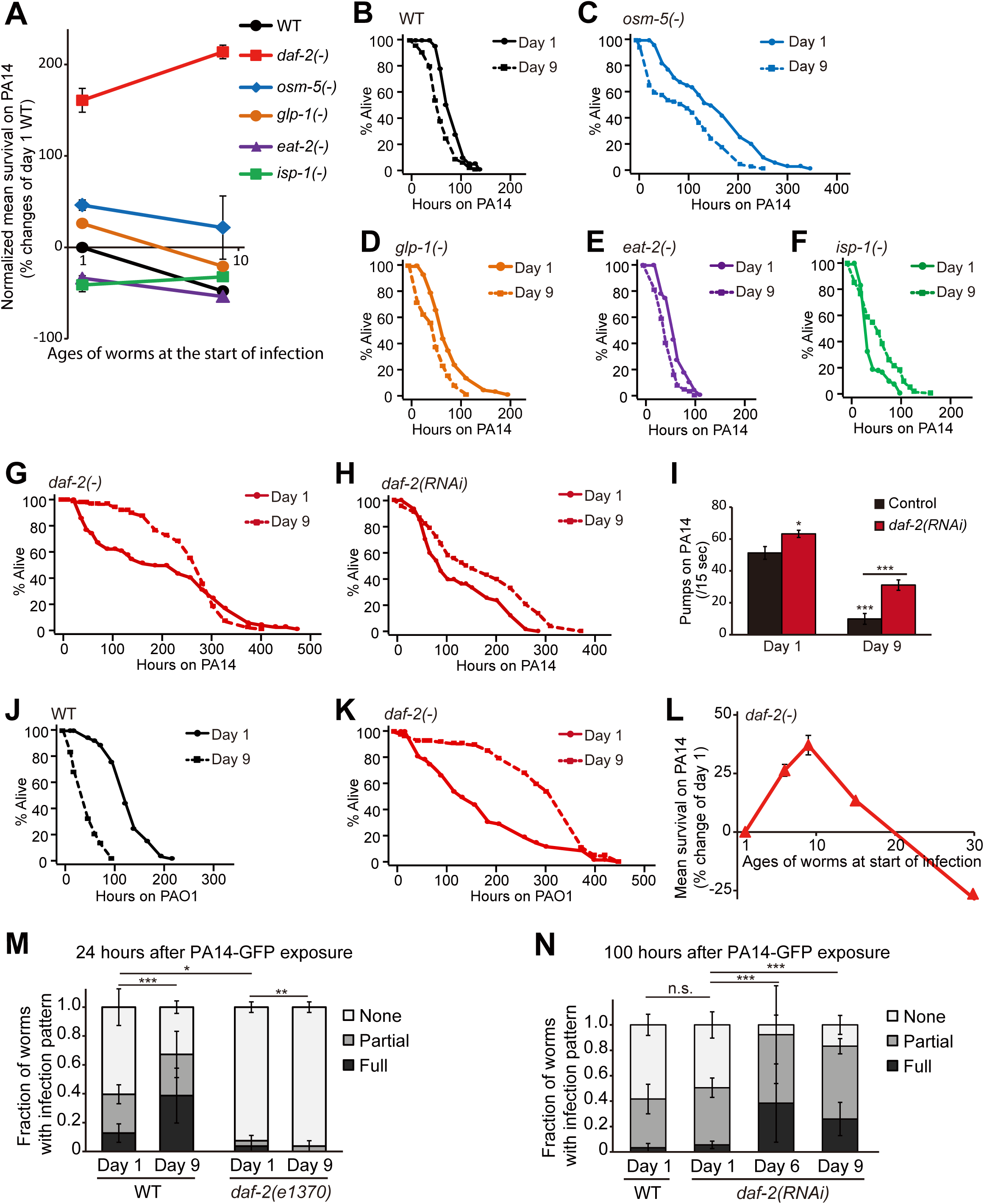
Old *daf-2(-)* animals display enhanced immune function against pathogenic *Pseudomonas aeruginosa*. (**A**) Normalized survival of wild-type (WT) and long-lived *daf-2(e1370)* [*daf-2(-)*], *osm-5(p813)* [*osm-5(-)*], and *glp-1(e2141)* [*glp-1(-)*], *eat-2(ad1116)* [*eat-2(-)*], and *isp-1(qm150)* [*isp-1(-)*] mutant animals transferred from normal food, *E. coli* OP50, to pathogenic *P. aeruginosa* (PA14) at day 1 or day 9 adulthood. WT worms displayed decreased survival on PA14 in an age-dependent manner. At all the ages that we tested, *daf-2(-)* mutants displayed enhanced survival on PA14. All survival assays except that with *glp-1(-)* animals were performed twice independently. Temperature-sensitive *glp-1(-)* mutants were maintained at 25°C until day 9 adulthood in the absence of 5-fluoro-2’-deoxyuridine (FUdR), a chemical inhibitor that prevents progeny from hatching by inhibiting DNA synthesis. (**B-H**) Shown are survival curves of WT (**B**), *osm-5(-)* (**C**), *glp-1(-)* (**D**), *eat-2(-)* (**E**), *isp-1(-)* (**F**), *daf-2(-)* (**G**), and *daf-2* RNAi-treated worms (**H**) transferred from *E. coli* to PA14 at day 1 and day 9 adulthoods. For four out of five trials, day 9 *daf-2(RNAi)* worms survived longer than day 1 *daf-2(RNAi)* worms on PA14. (**I**) Quantification of pharyngeal pumping rates of control RNAi- and *daf-2* RNAi-treated WT worms at day 1 and 9 adulthoods on PA14. See Figure S1A for *daf-2(-)* mutant pumping rate data. Error bars represent the standard error of mean (SEM) (two-tailed Student’s *t*-test, \**p<*0.05, \*\*\**p<*0.001). n = 20 from two independent experiments. (**J**-**K**) Survival curves of WT (**J**) and *daf-2(-)* (**K**) worms transferred from *E. coli* to *P. aeruginosa* (PAO1) at day 1 and day 9 adulthoods. (**L**) Normalized mean survival of day 1, 6, 9, 15 and 30 *daf-2(-)* mutants on PA14. Error bars represent the SEM. Two independent survival assays were performed except day 15 *daf-2(-)* animals (n=1). (**M**) Semi-quantification of PA14-GFP levels in WT and *daf-2(-)* worms at day 1 and day 9 adulthoods. *daf-2(-)* mutations suppressed the increased PA14-GFP accumulation with age (n ≥ 25 from 3 trials). *p* values were calculated by using chi-squared test (\**p* < 0.05, \*\**p* < 0.01, \*\*\**p* < 0.001). (**N**) *daf-2* RNAi-treated worms displayed an age-dependent increase in PA14-GFP accumulation (n ≥ 25 from 3 trials). See Figure S1C for representative images of worms. *p* values were calculated by using chi-squared test (\*\*\**p* < 0.001). See Table S2 for additional repeats and statistics analysis for survival data shown in Figure 1.

We then determined the duration of the enhanced PA14 resistance conferred by *daf-2* mutations, and found that *daf-2* mutant or *daf-2(RNAi)* animals maintained the PA14 resistance up to day 15 of adulthood, at a level that was comparable to that of young (day 1) adult animals (Figures 1L and S1B). Wild-type animals display an age-dependent increase in the intestinal accumulation of PA14, which is a sign of immunosenescence (Youngman et al., 2011). Therefore, we measured the accumulation of PA14 in the intestinal lumens of *daf-2* mutants and *daf-2(RNAi)* animals at different ages. We found that *daf-2* mutants displayed substantially reduced PA14 levels in the intestine in both young (Evans et al., 2008a) and old animals (Figure 1M). In contrast, we observed an age-dependent increase in the intestinal accumulation of PA14 in *daf-2* RNAi-treated animals (Figures 1N and S1C). As both old *daf-2* mutants and old *daf-2(RNAi)* worms survived longer on PA14 than did young animals (Figures 1G and 1H), genetic inhibition of *daf-2* appears to maintain immunocompetence in old age, at least in part, independently of PA14 intake and clearance.

### Temporal inhibition of *daf-2* specifically reverses immunosenescence

We next asked whether temporal genetic inhibition of *daf-2* was sufficient to enhance immunocompetence against PA14 at old age. We treated wild-type animals with control RNAi or *daf-2* RNAi starting at a middle age (day 4) and measured the survival at old age (day 8) (Figure 2A). Importantly, the temporal *daf-2* RNAi-treated old (day 8) worms displayed enhanced PA14 resistance compared with middle-aged (day 4) worms (Figure 2B). In contrast, the same population of control RNAi-treated worms displayed a substantial decline in survival upon PA14 infection at day 8 adulthood (Figure 2B). These data suggest that temporal inhibition of *daf-2* in wild-type worms during aging can rejuvenate immunity. In contrast to PA14 resistance, we found that the temporal *daf-2* RNAi treatment from day 4 did not reverse the age-dependent decline in resistance against oxidative stress or heat stress (Figures 2C and 2D). Thus, temporal inhibition of *daf-2* specifically enhances immunocompetence without increasing general stress resistance at old age.

**Figure 2.**
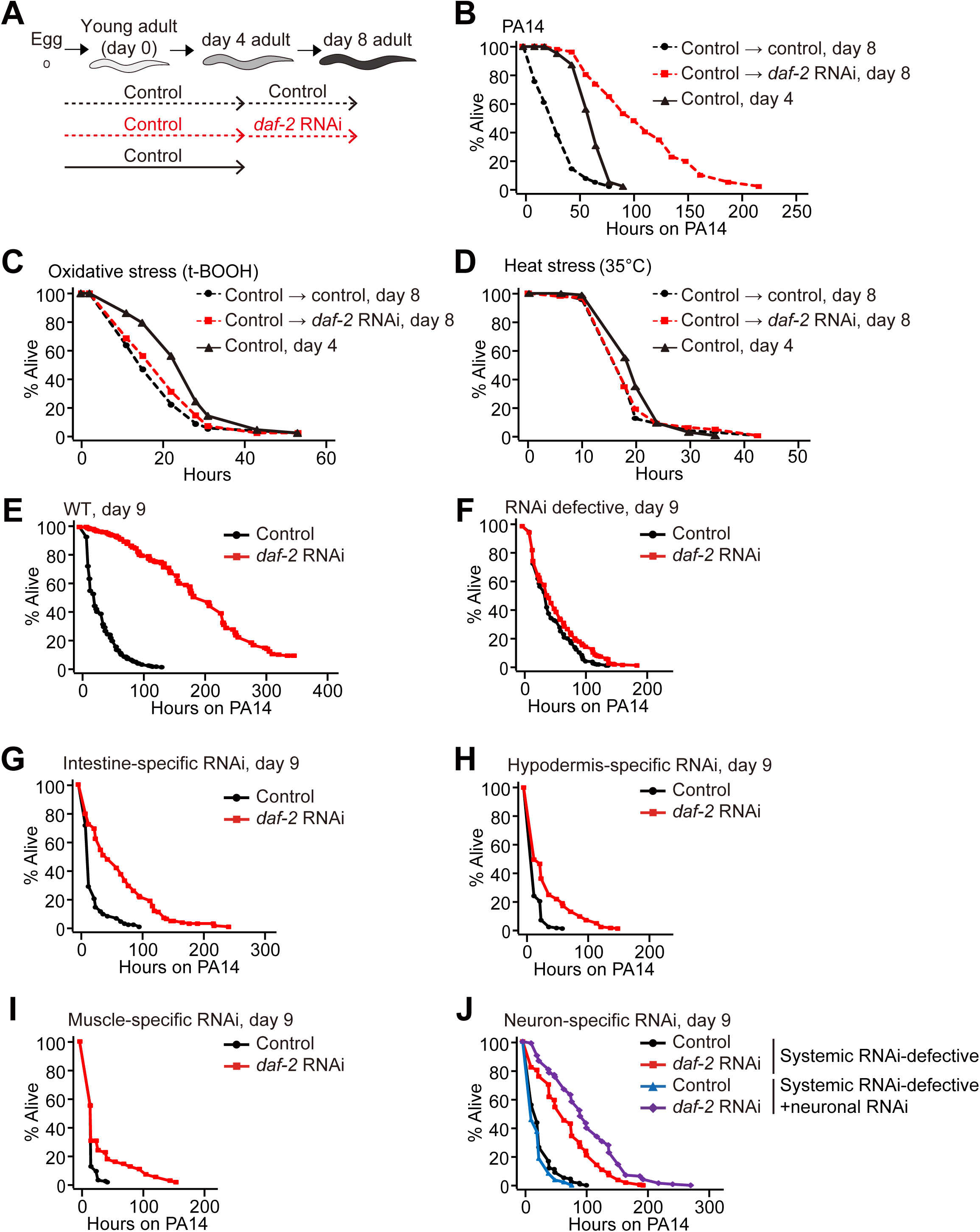
Temporal and spatial *daf-2* RNAi knockdown for modulation of reversed immunosenescence. (**A**) A schematic showing temporal *daf-2* RNAi experiments for PA14 and stress resistance assays shown in panels **B**-**D**. See materials and methods for details. (**B**-**D**) Temporal *daf-2* RNAi treatment from middle age (day 4 adulthood) increased PA14 resistance (3 trials) (**B**), but not oxidative (2 trials) (**C**) or heat stress resistance (1 trial) (**D**) in old (day 8 adulthood) worms. (**E**-**F**) *daf-2* RNAi delayed the immunosenescence of wild-type (WT) (**E**), but not that of RNAi-defective *rde-1(ne219)* [*rde-1(-)*] (**F**) animals at day 9 adulthood. (**G-I**) Intestine-(**G**), hypodermis-(**H**), and muscle-(**I**) specific *daf-2* RNAi increased PA14 resistance at day 9. (**J**) *daf-2* RNAi increased the PA14 resistance of systemic RNAi-defective *sid-1(pk3321)* [*sid-1(-)*] mutants that were used as a control for neuron-specific RNAi experiments. Neuron-specific *daf-2* RNAi treatment further increased PA14 resistance of *sid-1(-)* mutants at day 9 adulthood. Worms used for panels **E**-**J** were performed twice and pooled for survival curves. See Tables S3 and S4 for statistical analysis and additional for survival curves shown in Figure 2.

### Genetic inhibition of *daf-2* in the intestine, hypodermis, muscle, and neurons decreases susceptibility to pathogenic bacteria in old worms

We then assessed in which tissues DAF-2 affected immunosenescence by performing tissue-specific RNAi experiments (Calixto et al., 2010; Qadota et al., 2007). First, we showed that intestine-, hypodermis-, muscle-, and neuron-specific *daf-2* RNAi treatment significantly increased lifespan (Figures S2A-S2H), suggesting a role for reduced DAF-2 expression in life extension in these various tissues. We then found that intestine-, hypodermis-, muscle-, and neuron-specific *daf-2* RNAi treatment significantly enhanced PA14 resistance at old age (day 9), whereas it did not increase survival upon PA14 infection at young age (day 1) (Figures 2E-2J, S3A-S3F). These data suggest that reduced DAF-2 expression in the intestine, muscle, hypodermis, and neurons collectively contributes to the enhancement of immunocompetence at old age.

### *daf-2* mutation bypasses the requirement of p38 MAPK signaling for enhanced immunity at old age

Next, we sought to identify downstream factors that mediate the delayed immunosenescence in *daf-2* mutants. We first examined whether PMK-1/p38 MAPK, which is a major immune-regulatory factor and the critical determinant of immunosenescence in wild-type animals (Kim et al., 2002; Youngman et al., 2011), played a role in immunocompetence in old *daf-2* mutants. Consistent with the results of a previous study (Troemel et al., 2006), the enhanced PA14 resistance observed in young *daf-2* mutants was largely dependent on *pmk-1* (Figure S4A).

Unexpectedly, *pmk-1* was only partially required for the enhanced immunity of old (day 9) *daf-2(–)* mutants (Figure S4B). Overall, old (day 9 or 15) *daf-2(–); pmk-1(–)* double mutants displayed a substantially increased survival upon PA14 infection compared with young (day 1) animals (Figure 3A). Similarly, old *daf-2* mutants containing mutations in each of *nsy-1*/MAPKKK and *sek-1*/MAPKK, which act upstream of PMK-1 (reviewed in Irazoqui and Ausubel, 2010; Kim and Ewbank, 2018), survived longer on PA14 than young animals (Figures 3B, 3C, S4C-S4F). In addition, the levels of active phospho-PMK-1 proteins and the expression of *pmk-1*-regulated genes declined during aging in both wild-type and *daf-2* mutant worms (Figures 3D and 3E), consistent with a previous report (Wu et al., 2019). These data suggest that *daf-2* mutations bypass the functional requirement of MAPK-cascade components and act through other factors for maintaining immunity during aging.

**Figure 3.**
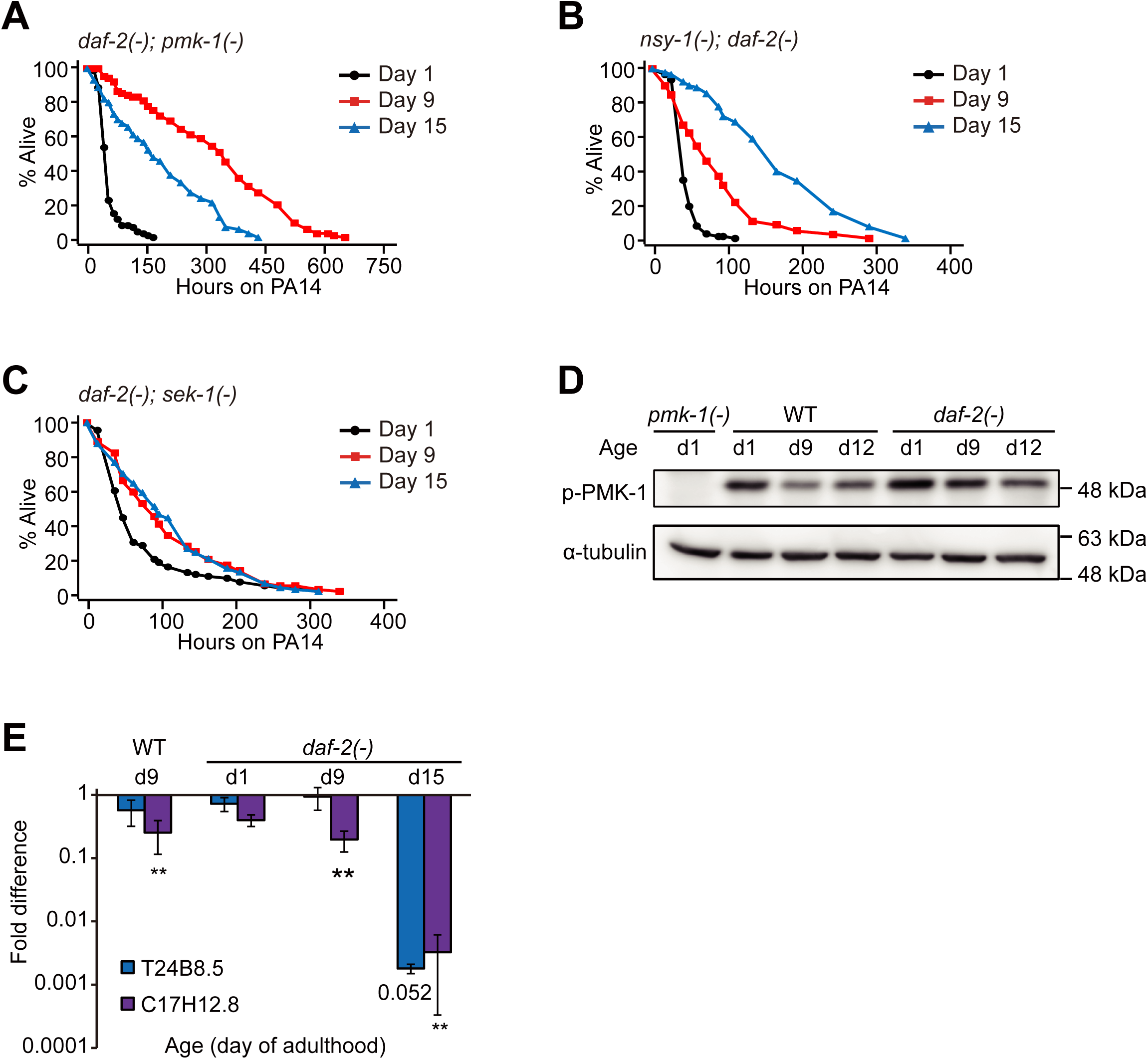
PMK-1 is dispensable for the delayed immunosenescence of *daf-2* mutants. (**A-C**) *daf-2* mutations delayed immunosenescence in *pmk-1(km25)* [*pmk-1(-)*] (**A**), *nsy-1(ok593)* [*nsy-1(-)*] (**B**) or *sek-1(km4)* [*sek-1(-)*] (**C**) mutant backgrounds. See Table S5 for statistical analysis and additional repeats. (**D**) Western blot analysis showed that the level of phospho-PMK-1 (p-PMK-1) was decreased with age in both wild-type (WT) and *daf-2(-)* mutant worms. Day (d) 1, 9, and 12 adult worms were used. The blot is a representative one out of three repeats that showed consistent results. α-tubulin was used as a loading control. (**E**) qRT– PCR analysis data indicate that the mRNA levels of two selected PMK-1-regulated genes, T24B8.5 and C17H12.8, were downregulated with age in both WT and *daf-2(-)* animals. Error bars represent SEM (two-tailed Student’s *t*-test., \*\**p*<0.01, n=3). *ama-1* was used as a normalization control. n (number of repeats) = 3 except T24B8.5 for *daf-2(-)* at day 15 (n=2).

### DAF-16/FOXO and HSF-1 are required for delayed immunosenescence in *daf-2* mutants

DAF-16/FOXO, HSF-1, and SKN-1/NRF are required for increased pathogen resistance in young *daf-2* mutants (Garsin et al., 2003; Miyata et al., 2008; Papp et al., 2012; Singh and Aballay, 2006, 2009 and Figures S5A, S5C and S5E); however, it remains unknown whether these transcription factors played a role in *daf-2(–)*-mediated delay in immunosenescence. Importantly, we found that *daf-16* mutations fully suppressed the PA14 resistance of *daf-2* mutants at all ages tested and unveiled immunosenescence in *daf-2* mutants (Figures 4A-4C, S5A and S5B). *hsf-1* RNAi also largely suppressed the enhanced immunocompetence of old *daf-2* mutants (Figures 4D-4F and S5D). In contrast, RNAi targeting *skn-1* did not affect the enhanced immunocompetence of old *daf-2* mutants (Figures 4G-4I and S5F). Together, these data suggest that DAF-16/FOXO and HSF-1 are responsible for drastically delayed immunosenescence in *daf-2* mutants.

**Figure 4.**
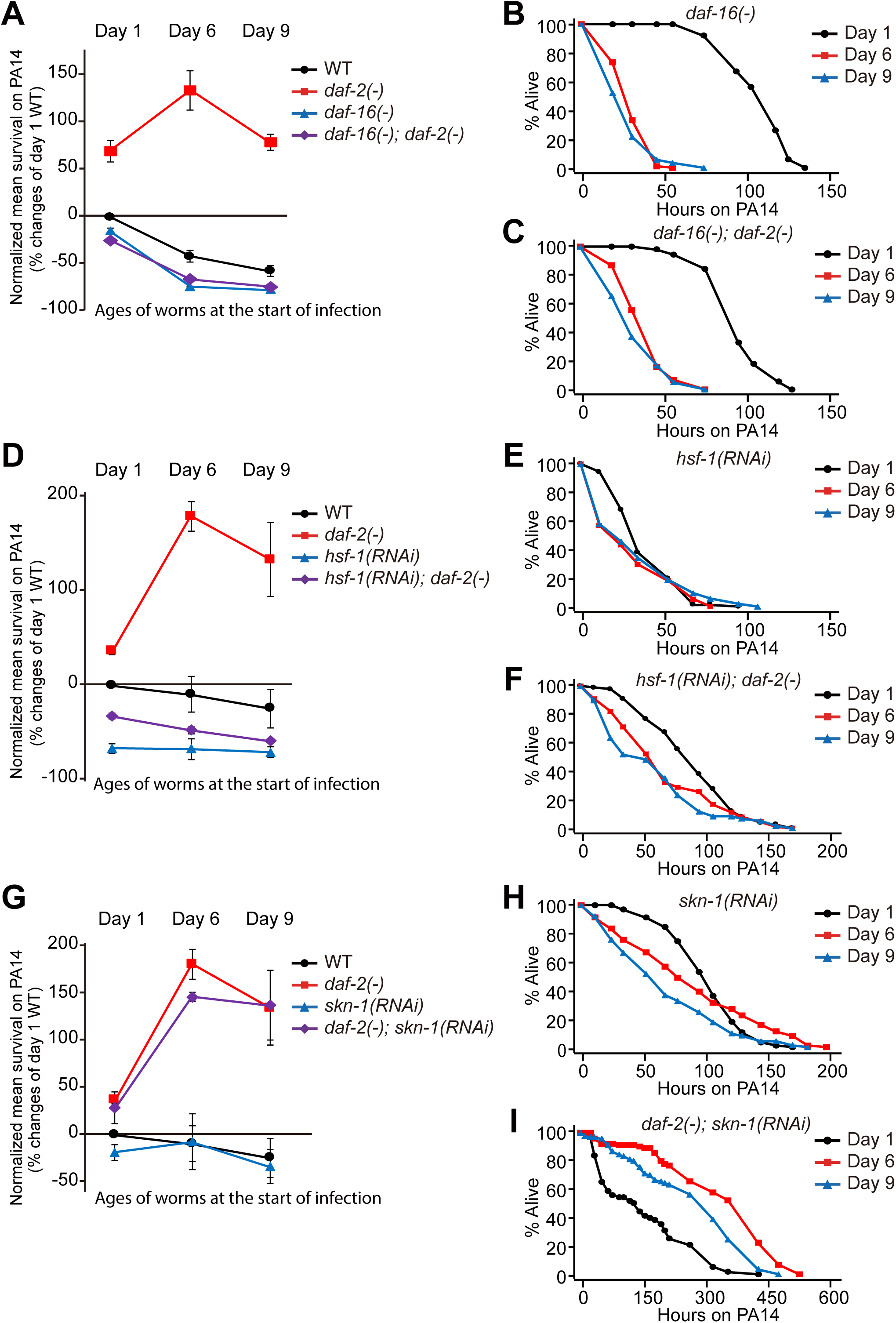
DAF-16 and HSF-1 are required for the immunocompetence in both young and old *daf-2* mutants. (**A**) Percent changes in mean survival upon PA14 infection among wild-type (WT), *daf-16(mu86)* [*daf-16(-)*], *daf-2(-)*, and *daf-16(-); daf-2(-)* animals at day 1, day 6, and day 9 adulthoods. (**B**) *daf-16(-)* mutants displayed an age-dependent increase in susceptibility to PA14. (**C**) *daf-2(-)* mutations also showed immunosenescence in a *daf-16(-)* mutant background. (**D**) Percent changes in mean survival of day 1, day 6, and day 9 *hsf-1* RNAi-treated WT and *daf-2(-)* animals on PA14. (**E**-**F**) *hsf-1* RNAi-treated WT (**E**) and *daf-2(-)* animals (**F**) displayed an age-dependent decline in PA14 resistance. (**G**) Percent changes in mean survival of *skn-1* RNAi-treated WT and *daf-2(-)* mutant animals on PA14. (**H**-**I**) *skn-1* RNAi treatment did not alter the immunosenescence of WT (**H**) or *daf-2(-)* animals (**I**). Error bars in panels **A, D**, and **G** represent SEM of mean survivals from two independent experiments. See Table S5 for statistical analysis and additional repeats for Figure 4.

### Age-associated gene expression changes are similar between wild-type and *daf-2* mutant worms

Next, we investigated which transcriptional changes were associated with and responsible for the enhanced immunocompetence observed in aged *daf-2* mutants. We conducted an mRNA sequencing (RNA seq) analysis using young (day 1) and old (day 9) *daf-2(–)* and wild-type animals. Hierarchical clustering indicated a clear separation of transcriptomes in accordance with ages and genotypes (Figure 5A), and a principal component analysis (PCA) revealed that the effect of age was larger than that of genotype (Figure S6A). Subsequently, we found that the age-dependent gene expression changes detected in wild-type worms displayed a positive correlation with those observed in *daf-2(–)* mutants (Figure 5B). We also showed that the mRNA levels of PMK-1 targets were decreased during aging in wild-type and *daf-2(–)* animals (Figure S6B), consistent with our Western blotting and qRT–PCR analysis (Figures 3D and 3E). In contrast, the mRNA levels of three selected DAF-16 targets, *mtl-1, hsp-16*.*2*, and *sod-3*, displayed age-dependent increases in wild-type worms, which tended to increase further in aged *daf-2(–)* mutants (Figures S6C-S6E). These data are consistent with our results showing the requirement of DAF-16/FOXO for the enhanced immunity observed in old *daf-2* mutants.

**Figure 5.**
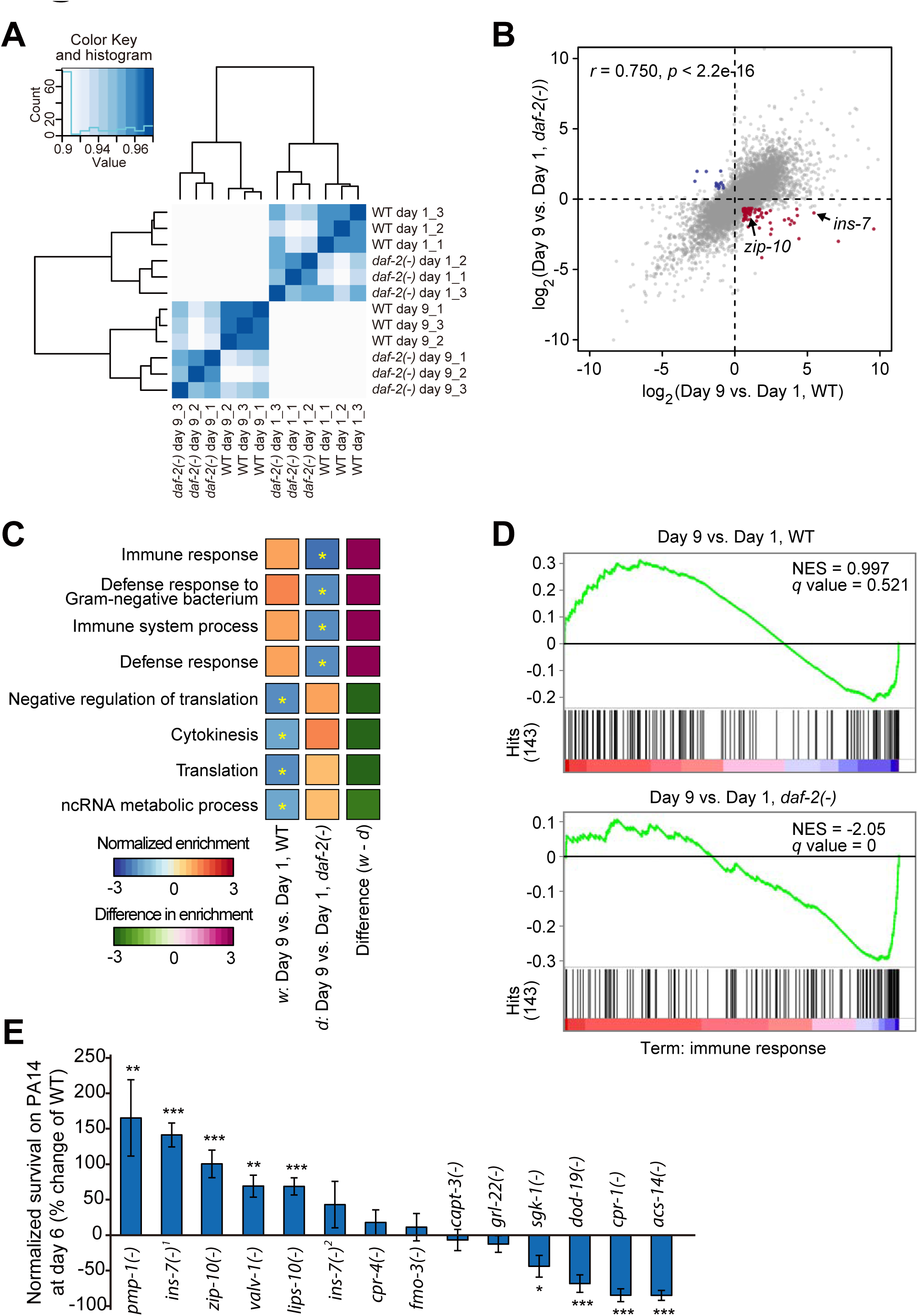
A subset of age-associated genes are differentially expressed in wild-type versus *daf-2* mutant worms. (**A**) A correlation matrix with hierarchical clustering of day 1 (young) and day 9 (old) adult wild-type (WT) and *daf-2(-)* animals (n=3). (**B**) A scatter plot showing gene expression changes conferred by *daf-2* mutation and aging. Genes whose expression was upregulated and downregulated with ages in WT but respectively downregulated and upregulated with ages in *daf-2(-)* animals (fold change > 1.5, BH-adjusted *p* value < 0.1) were shown in red and blue, respectively. *ins-7* and *zip-10* were indicated with arrows. (**C**) Results of GSEA showing terms significantly changed during aging in WT and *daf-2(-)* animals. Asterisks indicate terms significantly changed with significant *q* values (BH-adjusted *p* value < 0.1). *w*: differentially expressed genes (DEGs) with age in wild-type, *d*: DEGs with age in *daf-2(-)* animals. (**D**) Enrichment plots of GSEA show 143 ‘immune response’ genes were significantly downregulated during aging in *daf-2(-*) but not in WT animals. NES: normalized enriched score. *q* values were obtained by calculating the false discovery rate (FDR) corresponding to each NES. (**E**) Percent survival of animals that contain mutations in potential immunosenescence-regulating genes after PA14 infection for 48 hrs. Specific alleles that were used are as follows: *pmp-1(ok773)* [*pmp-1(ok773)*], *ins-7(tm1907)* [*ins-7(-)*^*1*^], *zip-10(ok3462)* [*zip-10(-)*], *valv-1(tm6726)* [*valv-1(-)*], *lips-10(tm7601)* [*lips-10(-)*], *ins-7(tm2001)* [*ins-7(-)*^*2*^], *cpr-4(ok3413)* [*cpr-4(-)*], *fmo-3(ok354)* [*fmo-3(-)*], *capt-3(ok1612)* [*capt-3(-)*], *grl-22(ok2704)* [*grl-22(-)*], *sgk-1(ok538)* [*sgk-1(-)*], *dod-19(ok2679)* [*dod-19*], *cpr-1(ok1344)* [*cpr-1(-)*], and *acs-14(ok3391)* [*acs-14(-)*]. We noticed that *ins-7(-)*^*1*^ elicited a stronger phenotype than *ins-7(-)*^*2*^ in this screen-type survival assays. However, our full survival assays indicated that both *ins-7(-)*^*1*^ and *ins-7(-)*^*2*^ mutations increased survival on PA14 at all ages that we tested (Figure 6E-G, Figure S7).

### Genetic inhibition of several genes whose expression is age-dependently altered enhances immunity in old age

We then asked which specific genes, whose expression was changed in an age-dependent manner, were responsible for the delayed immunosenescence of *daf-2(–)* mutants. Aged *daf-2(–)* mutants displayed increased immunity, which is the opposite of the impaired immunity observed in aged wild-type worms. Therefore, we focused our RNA seq analysis on genes that exhibited expression levels regulated in an opposite manner between wild-type and *daf-2(–)* worms during aging (Table S1). We found that the mRNA levels of 72 genes were increased with age in wild-type worms but were decreased with age in *daf-2(–)* animals (Figure 5B, red dots; fold change > 1.5; Benjamini and Hochberg (BH)-adjusted *p* value < 0.1 with 19,327 genes). Conversely, 14 genes exhibited expression levels that were decreased with age in wild-type worms and increased in *daf-2(–)* animals (Figure 5B, blue dots; fold change > 1.5; BH-adjusted *p* value < 0.1). A gene set enrichment analysis (GSEA) revealed that “immune/defense response” genes were significantly downregulated during aging in *daf-2(–)* worms but were marginally increased in wild-type animals (Figure 5C-5D, *q* value (BH-adjusted *p* value) < 0.1); thus, these genes may underlie the distinct immunity phenotypes observed between aged wild-type and aged *daf-2(–)* animals. We therefore performed immunosenescence assays using mutants that carried mutations in each of the 13 genes that were selected among the 72 candidate genes (Figure 5E), whose expression was specifically increased during wild-type aging and decreased during *daf-2(–)* aging. We found that the *pmp-1*/peroxisomal ABC transporter ABCD3, *ins-7*/an insulin-like peptide, *zip-10*/bZIP transcription factor, *valv-1*/Cysteine-Rich Intestinal Protein (CRIP), and *lips-10*/lipase loss-of-function mutants displayed enhanced immunocompetence at day 6 of adulthood compared with their wild-type counterparts (Figure 5E).

### An age-dependent decrease in *ins-7* expression contributes to enhanced immunocompetence in old *daf-2* mutants

Interestingly, among the genes that displayed an opposite age-dependent change in expression between *daf-2* mutants and wild-type animals, *ins-7* was negatively regulated by both DAF-16/FOXO (Figure S6F) and HSF-1 (Figure S6G). The *ins-7* gene encodes an agonist of DAF-2 that acts as a positive feedback regulator of IIS (Murphy et al., 2007; Murphy et al., 2003). Using qRT–PCR assays, we confirmed our RNA seq results by showing that the *ins-7* mRNA level was increased with age in wild-type worms, but not in *daf-2* mutants (Figures 6A and 6B). Consistent with previous reports (Lee et al., 2009; Murphy et al., 2007; Murphy et al., 2003), *ins-7* expression was increased by *daf-16(–)* (Figure 6C). In addition, *hsf-1* RNAi robustly elevated the level of the *ins-7* mRNA (Figure 6D). These data suggest that *ins-7* is downregulated by DAF-16/FOXO and HSF-1 in *daf-2* mutants but is upregulated during aging in wild-type worms. We then asked whether *ins-7* contributed to immunocompetence in old animals. Consistent with previous reports (Evans et al., 2008b; Kawli and Tan, 2008), *ins-7* mutation enhanced pathogen resistance at young age (day 1 adulthood) (Figures 6E and S7A). Notably, mutations in *ins-7* enhanced immunocompetence in adults at days 6 and 9 (Figures 6F-6G, S7B and S7C), indicating the immune-suppressive roles of INS-7 during aging. Taken together, these data suggest that the upregulation of DAF-16/FOXO and HSF-1 decreases *ins-7* expression and enhances immunocompetence further in old *daf-2* mutants.

**Figure 6.**
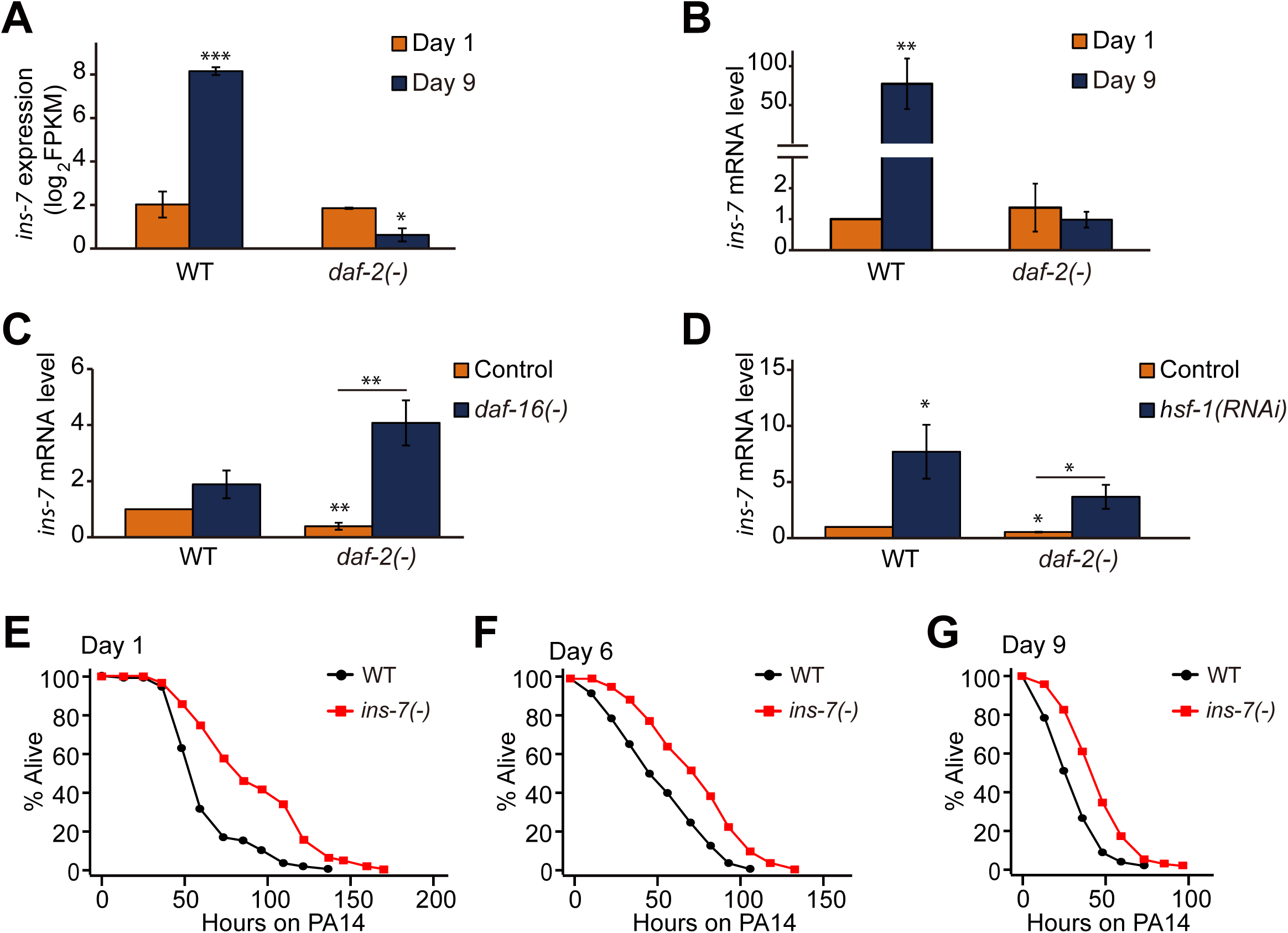
Down-regulation of *ins-7* contributes to enhanced immunocompetence in old *daf-2* mutants. (**A-B**) mRNA levels of *ins-7* in wild-type (WT) and *daf-2(-)* mutants at day 1 and day 9 adulthoods were measured by using RNA seq (**A**) and qRT-PCR (**B**). Error bars represent SEM (two-tailed Student’s *t*-test, n = 3). *ama-1* and *pmp-3* were used as normalization controls. (**C-D**) qRT-PCR analysis showed that *ins-7* mRNA levels were increased by *daf-16(-)* mutation (**C**) or *hsf-1* RNAi (**D**); the finding in panel **D** was first made by Coleen T. Murphy (unpublished) and was independently reproduced in this paper. Error bars represent SEM (two-tailed Student’s *t*-test, n = 4). *ama-1* was used as a normalization control. (**E**-**G**) *ins-7(tm1907)* [*ins-7(-)*^*1*^] mutation increased survival of day 1 (**E**), day 6 (**F**), and day 9 (**G**) adult worms on PA14. We obtained similar results by using *ins-7(tm2001)* [*ins-7(-)*^*2*^], another deletion mutant allele of *ins-7* (Figure S7). See Table S5 for additional repeats and statistical analysis.

### The ZIP-10/bZIP transcription factor acts as a positive regulator of *ins-7* for inducing immunosenescence

Next, we focused our functional characterization on the role of the ZIP-10/bZIP transcription factor in the regulation of immunosenescence with respect to INS-7 for the following reasons. First, a previous report indicated that ZIP-10 is a pathogen-responsive gene (Shapira et al., 2006). Second, the changes in the mRNA expression of *zip-10* were similar to those of *ins-7* in aged wild-type versus *daf-2* mutants (Figure 5B). Third, both *ins-7* and *zip-10* expression was increased by *daf-16* mutations and *hsf-1* RNAi in *daf-2* mutants (Figures 6C, 6D, 7A and 7B). Therefore, we wondered whether ZIP-10 acted together with INS-7 to regulate immunosenescence. We first showed that *zip-10* mutations suppressed the age-dependent increase in *ins-7p::GFP* expression (Figures 7C and 7D), suggesting that ZIP-10 induces *ins-7*. We then conducted a converse analysis using *isy-1(–)* mutants, which exhibit *zip-10* upregulation (Jiang et al., 2018). We found that *ins-7* was among the nine genes upregulated by both *isy-1(–)* and aging specifically in wild-type worms (Figure 7E). Therefore, ZIP-10 appears to upregulate *ins-7* during aging in wild-type worms, but not in *daf-2* mutants. We also showed that *zip-10* mutation increased the PA14 resistance at old age, but not at young age (Figures 7F and 7G). Conversely, *isy-1* mutations significantly reduced immune competence in old worms, but not in young worms (Figures 7H and 7I). Altogether these data suggest that *daf-2* mutations enhance immunocompetence in old age by downregulating ZIP-10; this decreases INS-7 expression, which normally elicits immunosenescence in wild-type worms.

**Figure 7.**
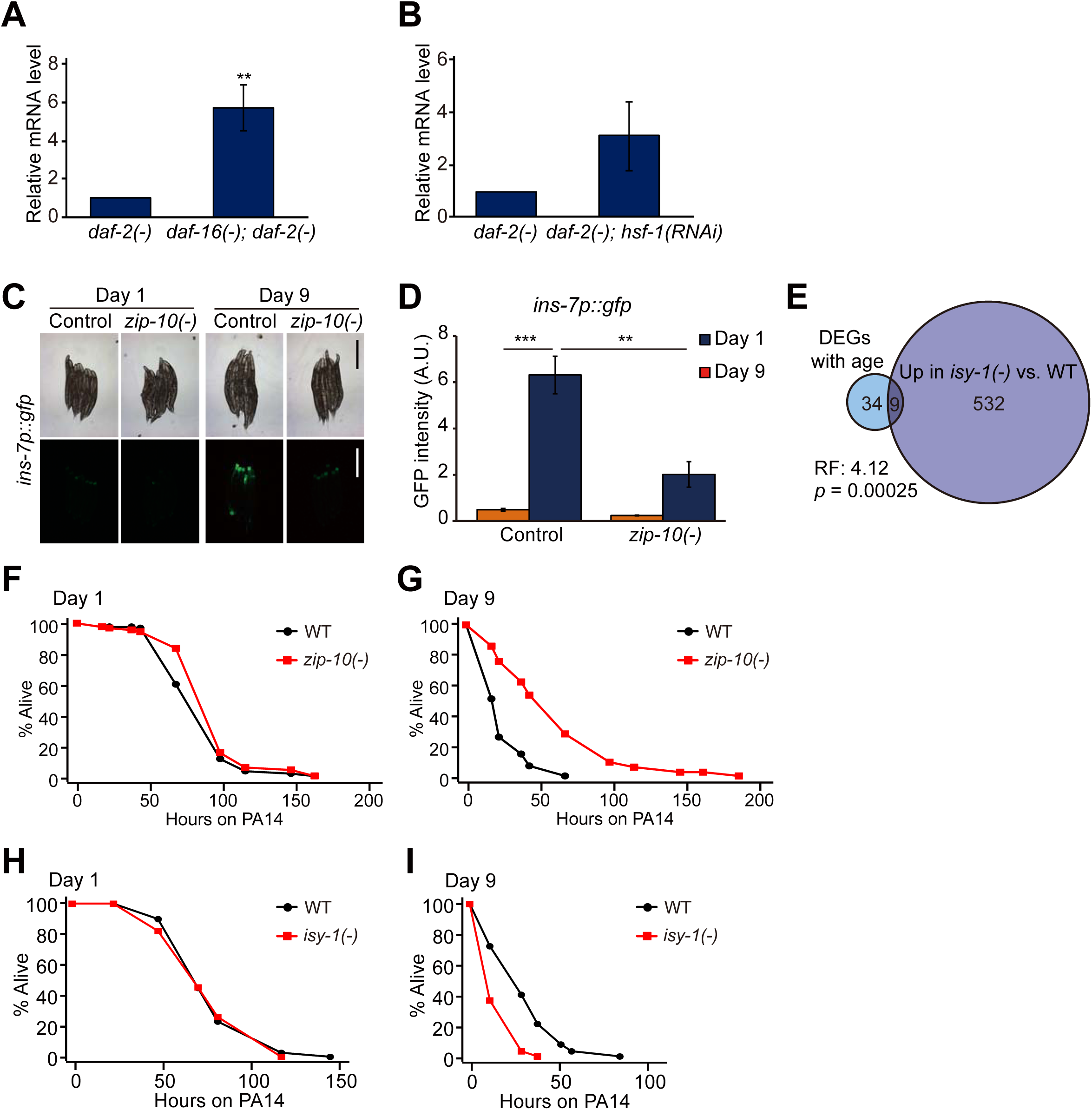
ZIP-10 positively regulates INS-7 during aging for inducing immunosenescence. (**A**-**B**) qRT-PCR analysis showed that the level of *zip-10* mRNA was increased by *daf-16(-)* mutation (**A**, \*\**p* < 0.01, n=7) or *hsf-1* RNAi (**B**, *p*= 0.181, n=3) in *daf-2(-)* animals. *ama-1* was used as a normalization control. Error bars represent SEM (two-tailed Student’s *t*-test). (**C**) Images of *ins-7p::gfp* [*ins-7p::gfp; unc-122p::dsred*] transgenic worms in wild-type (WT) and *zip-10(ok3462)* [*zip-10(-)*] mutant backgrounds at day 9 adulthood. Scale bar indicates 500 µm. (**D**) Quantification of the data shown in panel **C**. Error bars represent SEM (n ≥ 3, two-tailed Student’s *t-*tests, \*\**p* < 0.01, \*\*\**p* < 0.001). (**E**) Venn diagram represents a significant overlap between genes whose expression levels were oppositely regulated during aging between WT and *daf-2(-)* animals (labeled as “DEGs with age”) and genes whose expression was negatively regulated by ISY-1 (Jiang et al., 2018). RF: representation factor. *p* value was calculated with hypergeometric probability test. (**F**-**G**) *zip-10(-)* mutation did not affect the PA14 resistance of day 1 adults (**F**), but significantly increased the PA14 resistance of day 9 adults (**G**). (**H**-**I**) *isy-1(dma50)* [*isy-1(-)*] mutation did not affect the susceptibility of day 1 adult worms against PA14 (**H**), but significantly decreased the PA14 resistance in day 9 adult worms (**I**). See Table S5 for additional repeats and statistics.

## Discussion

### Reduced IIS appears to reverse immunosenescence *via* a positive endocrine feedback loop

Despite the close association between aging and immunity, whether and how longevity interventions affect immunosenescence remains largely unexplored. Here, we determined a relationship between longevity and immunosenescence using *C. elegans*. We found that reduction of IIS greatly delayed and even rejuvenated immunity at some point during aging. This retarded immunosenescence conferred by inhibition of the DAF-2/insulin/IGF-1 receptor was dependent on DAF-16/FOXO and HSF-1, whereas PMK-1/p38 MAPK signaling and SKN-1/NRF were dispensable. These data suggest that the activation of DAF-16/FOXO and/or HSF-1 is sufficient to reverse immunosenescence, whereas the activation of p38 MAPK is not a limiting factor for delaying immunosenescence. We also showed that *daf-2* mutations suppressed the age-dependent upregulation of the ZIP-10/bZIP transcription factor, which led to the downregulation of INS-7 and boosted immunity. Because INS-7 is an agonist of DAF-2 (Murphy et al., 2003), this in turn appears to act as a positive feedback circuit to enhance immunity further in old *daf-2* mutants. We propose that *daf-2* mutations can uncouple aging and the age-dependent declines in immune function by increasing the activities of DAF-16/FOXO and HSF-1, but not those of p38 MAPK signaling, *via* a positive-feedback mechanism during aging.

### Pharyngeal pumping does not appear to be a major cause of delayed immunosenescence in *daf-2(–)* animals

PA14 colonizes the *C. elegans* intestinal lumen, and exposure to PA14 ceases pharyngeal pumping in *C. elegans* (Tan et al., 1999). We wondered whether the reduction of pharyngeal pumping with age caused increased pathogen resistance in *daf-2(–)* mutants. Although animals with reduced *daf-2* functions displayed age-dependent decreases in pharyngeal pumping after infection with the pathogen (Figure 1I and S1A), several lines of evidence indicated that pharyngeal pumping is not the main cause of the increased immunocompetence observed in old *daf-2(–)* mutants. First, aged wild-type worms displayed a reduced pumping rate but showed increased susceptibility to PA14. Second, *eat-2(–)* mutants with defective pharyngeal pumping displayed an age-dependent reduction in pathogen resistance. Third, *daf-2* RNAi treatment significantly improved pharyngeal pumping and survival upon PA14 infection compared with wild-type worms. Fourth, *daf-2* RNAi treatment in post-reproductive, middle-aged, wild-type animals was sufficient to enhance immunocompetence in old age. Overall, these data suggest that delayed immunosenescence can be achieved without defects in feeding rates.

### The intestine and neurons are important tissues for *daf-2(–)*-mediated delayed immunosenescence, enhanced immunity, and longevity

Previous reports have shown that the expression of DAF-16/FOXO and HSF-1 in various tissues contributes to the longevity of *daf-2* mutants (Douglas et al., 2015; Libina et al., 2003; Morley and Morimoto, 2004; Son et al., 2018; Zhang et al., 2013). In addition, tissue-specific transgenic rescue analyses indicate that the loss of *daf-2* function in neurons is required for longevity (Wolkow et al., 2000). However, the specific tissues in which reduced DAF-2 function is sufficient for extending lifespan remain unknown. Here, we demonstrated that reducing *daf-2* in all tissues tested (intestine, hypodermis, muscle, and neurons) contributed to longevity (Figure S2). Importantly, we also showed that intestine-, hypodermis-, muscle-, and neuron-specific *daf-2* RNAi enhanced immunocompetence at old age (day 9 adulthood). Our results are consistent with those of previous reports, which showed that the intestine, hypodermis, and neurons are tissues important for the regulation of immunity (Irazoqui and Ausubel, 2010; Kim and Ewbank, 2018). The intestine and neurons were shown to be the main tissues that express *ins-7* (Chen et al., 2013; Matsunaga et al., 2016; Murphy et al., 2007; Murphy et al., 2003). Thus, our data raise the possibility that the expression and subsequent secretion of INS-7 in the intestine and neurons act in a cell-non-autonomous manner on DAF-2 in multiple tissues, thus modulating immunosenescence and longevity.

### Downregulation of the INS-7/insulin-like peptide may delay immunosenescence

The secretion of INS-7 by the nervous system negatively regulates immunity-related factors in intestinal cells *via* the DAF-16/FOXO transcription factor (Evans et al., 2008b; Kawli and Tan, 2008; Murphy et al., 2003). Our data suggest that the age-dependent increase in *ins-7* expression is associated with immunosenescence in *C. elegans*. Consistent with previous reports (Murphy et al., 2007; Murphy et al., 2003), we showed that *ins-7* expression increased with age in wild-type worms. Notably, we found that *ins-7* expression decreased slightly with age in *daf-2(–)* mutants. Moreover, we demonstrated that genetic depletion of *ins-7* enhanced pathogen resistance in old worms. Overall, these data suggest that inhibition of INS-7 during aging is an effective way of reversing immunity in old age.

### IIS may play roles in mammalian immunosenescence

Previous studies have shown that IIS regulates innate immunity in various other species. FOXO, which is a conserved transcription factor that acts downstream of IIS, promotes innate immunity directly in *Drosophila* and mammalian cells (Becker et al., 2010). Reduced IGF-1 signaling promotes the renewal of hematopoietic stem cells (HSCs), which are crucial for the generation of cells important for proper immune functions (Cheng et al., 2014). In addition, inhibition of target of rapamycin, which intersects IIS, increases vaccination response in elderly people and enhances the self-renewal of HSCs in mice (Chen et al., 2009; Mannick et al., 2014). In the current study, we identified systemic positive feedback mechanisms by which IIS rejuvenates immunity in *C. elegans*. Because the functions of IIS in various physiological processes, including aging and immunity, are conserved across phyla, the findings reported here for *C. elegans* may lead to the development of therapeutic strategies against immunosenescence in humans.

## Materials and Methods

### *C. elegans* strains and maintenance

All strains were maintained as previously described (Stiernagle, 2006). Some strains were obtained from Caenorhabditis Genetics Center, which is funded by the NIH National Center for Resources (p40 OD010440), or National Bio-Resource Project (NBRP). Strains used in this study are as follows: N2 wild-type, CF1041 *daf-2(e1370) III*, CF2553 *osm-5(p813) X*, CF1903 *glp-1(e2141) III*, CF2172 *isp-1(qm150) IV*, IJ173 *eat-2(ad1116) II* obtained by outcrossing DA1116 four times to Lee-laboratory N2, WM27 *rde-1(ne219) V*, NR222 *rde-1(ne219) V; kzls9[lin-26p::nls::gfp, lin-26p::rde-1; rol-6D]*, NR350 *rde-1(ne219) V; kzls20[hlh-1p::rde-1; sur-5p::nls::gfp]*, VP303 *rde-1(ne219) V; kbls7[nhx-2p::rde-1; rol-6D]*, NL3321 *sid-1(pk3321) V*, TU3401 *sid-1(pk3321) V; uIs69[myo-2p::mCherry; unc119p::sid-1]*, IJ130 *pmk-1(km25) IV* obtained by outcrossing KU25 four times to Lee-laboratory N2, IJ1147 *daf-2(e1370) III; pmk-1(km25) IV*, IJ128 *sek-1(km4) X* obtained by outcrossing KU4 four times to Lee-laboratory N2, IJ1299 *daf-2(e1370) III; sek-1(km4) X*, IJ131 *nsy-1(ok593) II* obtained by outcrossing VC390 four times to Lee-laboratory N2, IJ1298 *nsy-1(ok593) II; daf-2(e1370) III*, CF1042 *daf-16(mu86) I*, CF1085 *daf-16(mu86) I; daf-2(e1370) III*, IJ1733 *ins-7(tm1907)* obtained by outcrossing FX01907 four times to Lee-laboratory N2, IJ1010 *ins-7(tm2001)* obtained by outcrossing QL126 four times to Lee-laboratory N2, ZC1436 *yxIs13[ins-7p::gfp(70ng/ul); unc-122p::dsred]* a gift from Yun Zhang laboratory, IJ1873 *daf-2(e1370) III; yxIs13[ins-7p::gfp(70ng/ul); unc-122p::dsred]*. IJ1871 *zip-10(ok3462)* obtained by outcrossing RB2499 five times to Lee-laboratory N2, IJ1919 *isy-1(dma50)* obtained by outcrossing DMS175, a gift from Dengke Ma laboratory. RB908 *pmp-1(ok773)*, RB1262 *cpr-1(ok1344)*, RB1415 *capt-3(ok1612)*, RB2452 *acs-14(ok3391)*, RB2473 *cpr-4(ok3413)*, RB2499 *zip-10(ok3462)*, RB686 *fmo-3(ok354)*, FX06726 *valv-1(tm6726)*, FX07601 *lips-10(tm7601)*, FX01907 *ins-7(tm1907)*, CF2078 *sgk-1(ok538)*, VC2249 *dod-19(ok2679)*, and VC2200 *grl-22(ok2704)*. All strains except *daf-2(e1370); sek-1(km4)* were maintained on NGM plates seeded with *Escherichia coli* OP50 strain as a food source at 20°C. *daf-2(e1370); sek-1(km4)* was maintained and synchronized at 15°C because of a semi-sterility phenotype at 20°C.

### Immunosenescence assays

Immunosenescence assays were performed as described previously (Youngman et al., 2011) with minor modifications. Gravid adult worms were allowed to lay eggs overnight to synchronize progeny on nematode growth media (NGM) plates seeded with *E. coli* OP50 or RNAi bacteria. When the progeny reached L4 or young adult stage (day 0), the worms were transferred onto NGM plates treated with 50 μM of 5-fluoro-2’-deoxyuridine (FUdR, Sigma, St. Louis, MO, USA). Worms were maintained on these plates at 20°C until they were used for experiments at day 1, 3, 6, 9, 12, or 15 of adulthood. For the immunosenescence assays using temperature-sensitive *glp-1(e2141)* mutants, N2 and *glp-1(e2141)* worms were synchronized by allowing gravid adults to lay eggs on *E. coli* OP50-seeded plates for 12 hrs at 20°C and the progeny developed at 25°C until the worms reached L4 stage. For the immunosenescence assays using *daf-2(e1370); sek-1(km4)* mutants, which were semi-sterile at 20°C, worms were first cultured at 15°C for obtaining a sufficient number of animals and transferred to 20°C for allowing to age, when they reached adulthood. Worms were transferred every other day until N2 worms stopped producing progeny. Contaminated or *E. coli*-depleted plates over time were discarded. PA14 standard slow killing assays were performed as previously described (Jeong et al., 2017). Briefly, 5 μl of overnight *P. aeruginosa* liquid culture was seeded onto the center of 0.35% peptone NGM solid media. Plates were incubated at 37°C for 24 hrs and subsequently at room temperature for over 8 hrs before use. Different ages of worms were simultaneously infected with PA14 on plates containing 50 μM of FUdR at 25°C. After infection (time = 0 on survival curves), worms were scored more than once a day by gently prodding with a platinum wire to determine life or death. Statistical analysis of survival data was performed by using OASIS (http://sbi.postech.ac.kr/oasis), which calculates *p* values using log-rank (Mantel-Cox method) test (Yang et al., 2011). For Figure 5E, worms were synchronized on OP50-seeded NGM plates and transferred to 50 μM FUdR-treated NGM plates when they became young (day 0) adults. After 6 days, worms were infected with PA14. After 48 hrs of infection, live and dead worms were counted.

### Temporal *daf-2* RNAi experiments

Wild-type worms were synchronized on control RNAi-treated plates and transferred onto new control RNAi plates when they became day 0 adults. Day 4 adult worms were transferred to control or *daf-2* RNAi bacteria-containing plates, respectively, and maintained at 20°C for 4 days. Day 8 adult worms were moved to plates with PA14 (Figure 2A), or OP50 plates with 7.5 mM t-BOOH (tert-Butyl hydroperoxide solution, Sigma, St Louis, MO, USA) (Figure 2B), or placed in a 35°C incubator (Figure 2C) and live worms were scored over time. OP50-seeded NGM plates were used for the heat and oxidative stress resistance assays with temporal *daf-2* RNAi-treated animals to avoid continuous RNAi induction and to match the RNAi conditions used for PA14 survival assays. Worms that reached young (day 0) adulthood were treated with 50 μM FUdR until finishing the survival assays. Statistical analysis of survival data was performed by using OASIS (http://sbi.postech.ac.kr/oasis), which calculates *p* values using log-rank (Mantel-Cox method) test (Yang et al., 2011).

### RNAi induction

Each RNAi clone-expressing HT115 bacteria were cultured in Luria broth (LB) containing 50 μg/ml ampicillin (USB, Santa Clara, CA, USA) overnight at 37°C. One hundred microliter of RNAi culture was seeded onto NGM containing 50 μg/ml ampicillin, and incubated overnight at 37°C. One mM isopropylthiogalactoside (IPTG, Gold biotechnology, St Louis, MO, USA) was added and incubated at room temperature for over 24 hrs before use.

### Lifespan assays

Lifespan assays were conducted as previously described (Son et al., 2017). Young (day 0) adult worms that were grown on control RNAi and *daf-2* RNAi plates were transferred onto 5 μM FUdR-containing control and *daf-2* RNAi plates, respectively. The animals that did not respond to gentle touch were considered as dead. The animals that crawled off the plates, missing, burrowed, or displayed vulval protrusion or internal hatching were censored but included in the subsequent statistical analysis. Statistical analysis of lifespan data was performed by using OASIS (http://sbi.postech.ac.kr/oasis), which calculates *p* values using log-rank (Mantel-Cox method) test (Yang et al., 2011).

### Intestinal PA14-GFP accumulation assays

Intestinal PA14-GFP accumlation assays were performed as previously described (Jeong et al., 2017; Kim et al., 2002), with minor modifications. Day 1 and day 9 adult N2 and *daf-2(e1370)* worms were infected with PA14 that expresses GFP (PA14-GFP) for 24 hrs and subsequently imaged using a Zeiss Axio Scope A1 compound microscope (Zeiss Corporation, Jena, Germany). For control and *daf-2* RNAi treatments, worms were infected with PA14-GFP for 100 hrs.

### Measurements of pharyngeal pumping

Pharyngeal pumping rates were determined as previously described (Cao and Aballay, 2016). PA14 bacterial lawns were prepared as described above. Aged animals were transferred onto PA14 lawns, and incubated at 25°C. The number of contractions of the terminal bulb was counted for 15 sec. The pumping rates for ten adults were measured for each condition.

### Microscopy

Worms were prepared at indicated ages and transferred on a 2% agarose pad with 100 mM sodium azide (DAEJUNG, Siheung, South Korea) or 2 mM levamisole (tetramisole; Sigma). The images of worms were captured by using an AxioCam HRc CCD digital camera (Zeiss Corporation, Jena, Germany) with a Zeiss Axio Scope A1 compound microscope (Zeiss Corporation, Jena, Germany). ImageJ (https://imagej.nih.gov/ij/) was used for the quantification of fluorescent images.

### RNA extraction and quantitative RT-PCR

RNA extraction and qRT-PCR were performed as previously described with minor modifications (Son et al., 2017). When worms that were fed with OP50 or RNAi bacteria reached L4 or young adult stage (day 0), 50 μM FUdR was supplemented. After 1 or 9 days, worms were harvested by washing with M9 buffer 2-3 times to remove residual bacteria. Plates with bactrial or fungal contamination, or with starved worms because of depletion of foods, were discarded. If OP50 was almost depleted, soaked freeze-dried OP50 (LabTIE, Rosmalen, Netherlands) was added onto the plates to prevent starvation. Total RNA in the animals was extracted using RNAiso plus (Takara, Seta, Kyoto, Japan) and cDNA was synthesized by ImProm-II™ Reverse Transcriptase kit (Promega, Madison, WI, USA). qRT-PCR was performed by using StepOne Real-Time PCR System (Applied Biosystems, Foster City, CA, USA) with SYBR green dye (Applied Biosystems, Foster City, CA, USA). Comparative C^T^ method was used for quantitative PCR analysis. Two technical repeats were averaged per biological sample and analyzed for all data sets. For all biological data sets, *ama-1* (an RNA polymerase II large subunit), and/or *pmp-3* (a peroximal membrane protein) were used as reference genes for normalization (Hoogewijs et al., 2008; Taki and Zhang, 2013)). Sequences of primers used for qRT-PCR are as follows. *ama-1*: 5’-TGGAACTCTGGAGTCACACC and 5’-CATCCTCCTTCATTGAACGG; *pmp-3*: 5’-GTTCCCGTGTTC ATCACTCAT and 5’-ACACCGTCGAGA AGCTGTAGA; *T24B8*.*5*: 5’-TGTTAGACAATGCCATGATGAA and 5’-ATTGGCTGTGCAGTTGTACC; *C17H12*.*8*: 5’-GAACAATAGTGTCAAGCCGATCTGC and 5’-TTCTGAATGATGAATGCATGTTTAC; *mtl-1*: 5’-GACTGCTGAAATTAAGAAATCATG and 5’-GTCTCCACTGCATTCACATTTGTC; *hsp-16*.*1/11*: 5’-CTCATGAGAGATATGGCTCAG and 5’-CATTGTTAACAATCTCAGAAG; *ins-7*: 5’-GTGGAAGAAGAATACATTCG and 5’-CTTCAGTATTCGATTCGCATG. *zip-10*: 5’-TCGAGATGCTCTTCAACTG and 5’-CTAACTGCTTGCCGGAG

### mRNA library preparation for RNA seq analysis and data acquisition

Synchronized N2 and *daf-2(e1370)* animals were treated with 50 μM FUdR when worms reached L4 or young adult stage on OP50. At indicated ages (day 1 and day 9 adulthoods), worms were harvested with M9 buffer, and washed 2-3 times, and frozen in liquid nitrogen. Three biological repeats of the samples were used for all conditions. For RNA seq analysis using *hsf-1* RNAi, *daf-2(e1370)* mutants were synchronized on control or *hsf-1* RNAi bacterial plates. Day 1 adult *daf-2(e1370)* animals on control or *hsf-1* RNAi plates were harvested with M9 buffer. Two biological repeats of samples were used. RNA extraction for RNA seq samples was carried out as described above (see RNA extraction section). TruSeq (unstranded) mRNA libraries (Illumina, CA, USA) were constructed and paired-end sequencing of Illumina HiSeq2500 was performed by Macrogen Inc. (Seoul, South Korea).

### RNA seq analysis

Sequencing pairs were aligned to the *C. elegans* genome WBcel235 (ce11) and Ensembl transcriptome (release 97) by using STAR (v.2.7.0e). Aligned pairs on genes or transcripts were quantified by using RSEM (v.1.3.1). Detailed parameters of alignment and quantification were calculated by following guidelines of ENCODE long RNA seq processing pipeline. Differentially expressed genes (fold change > 1.5 and adjusted *p* value < 0.1) were identified by using DESeq2 (v.1.22.2). Wald test *p* values are adjusted for multiple testing using the procedure of BH. Gene ontology (GO) terms generally changed in a certain comparison were identified by GSEA (v.4.0.1) using read counts of all expressed genes with a parameter ‘Permutation type: gene_set’. Terms with *q* value (BH-adjusted *p* value) less than 0.1 were considered as significantly changed. R (v.3.6.1, http://www.r-project.org) was used for plotting results. Raw data and processed data are available in Gene Expression Omnibus (https://www.ncbi.nlm.nih.gov/geo, GSE137861).

### Western blot assays

Synchronized N2, *daf-2(e1370)* and *pmk-1(km25)* animals were treated with 50 μM FUdR when worms reached L4 or young adult stage (day 0) on OP50. At indicated ages (day 1, 9, 12 and 15), worms were harvested with M9 buffer, and washed 2-3 times, and frozen in liquid nitrogen with 2X SDS sample buffer. Western blot assays were conducted as described previously (Jeong et al., 2017).

## Acknowledgments

We thank Drs. Dennis H. Kim, You-Hee Cho, Emily Troemel, Yun Zhang, Dengke Ma and the Caenorhabditis Genetics Center, for providing some bacteria and *C. elegans* strains. We also thank all Lee laboratory members and Dr. Kyuhyung Kim for help and discussion. This study is supported by the Korean Government (MSIT) through the National Research Foundation of Korea (NRF) (NRF-2016R1E1A1A01941152 and NRF-2019R1A3B2067745) to S-J.V.L. C.T.M. is the Director of the Glenn Center for Aging Research at Princeton University.

## Supplemental Figure Legends

**Figure S1. Pharyngeal pumping rates, age-dependent survival on PA14 and the accumulation of PA14 of animals with genetic inhibition of *daf-2***. (**A**) Pharyngeal pumping rates of day 1, 6 and 9 adult wild-type (WT) and *daf-2(e1370)* [*daf-2(-)*] animals on PA14. (**B**) Normalized mean survival of day 1, 9, 15, and 30 adult *daf-2* RNAi-treated worms on PA14. Error bars represent the SEM from five independent survival assay data for day 1 and day 9 old worms, and two independent survival assay data for day 15 and 30 old worms. (**C**) *daf-2* RNAi-treated worms displayed an age-dependent increase in PA14-GFP accumulation. Shown are representative images of worms that were pre-treated with *daf-2* RNAi, after PA14-GFP exposure for 100 hrs. * indicates PA14-GFP (scale bar: 100 μm). See Figure 1N for Semi-quantification of PA14-GFP levels shown in panel **C**.

**Figure S2. Genetic inhibition of *daf-2* in the intestine, hypodermis and neurons increases lifespan**. (**A**-**C**) *daf-2* RNAi increased the lifespan of wild-type (WT, N2) (**A**), but did not influence the lifespan of RNAi-defective *rde-1(ne219)* [*rde-1(-)*] (**B**) animals. (**C**-**E**) Intestine-(**C**), hypodermis-(**D**), and muscle-(**E**) specific *daf-2* RNAi significantly increased lifespan. (**F**-**G**) *daf-2* RNAi did not increase the lifespan of systemic RNAi-defective *sid-1(pk3321)* [*sid-1(-)*] mutants (**F**), while increasing the lifespan of neuron-specific RNAi strain (**G**). *rde-1(-)* mutant animals that expressed *rde-1* under the control of an intestine-specific *nhx-2* promoter, a hypodermis-specific *lin-26* promoter, or a muscle-specific *hlh-1* promoter, and *sid-1(-)* mutant animals that expressed *sid-1* driven by a neuron-specific *unc-119* promoter were used for tissue-specific RNAi experiments. (**H**) Normalized mean lifespan. Error bars represent the standard error of mean (SEM) (two-tailed Student’s *t*-test., \**p<*0.05, \*\**p<*0.01). Two independent lifespan assays were performed. See Table S4 for additional repeats and statistics.

**Figure S3. Inhibition of *daf-2* in single tissues did not increase the PA14 resistance of young (day 1) adult worms**. (**A**-**B**) *daf-2* RNAi significantly increased the survival of young (day 1) wild-type (WT) animals on PA14 (**A**), but not that of RNAi-defective *rde-1(-)* mutants (**B**). (**C**-**F**) Intestine-(**C**), hypodermis-(**D**), muscle-(**E**) or neuron-(**F**) specific *daf-2* RNAi did not increase the survival of young (day 1) adult worms. RNAi-defective *rde-1(ne219)* [*rde-1(-)*] mutant animals that expressed *rde-1* under the control of an intestine-specific *nhx-2* promoter, a hypodermis-specific *lin-26* promoter or a muscle-specific *hlh-1* promoter were used for tissue-specific RNAi experiments. Systemic RNAi-defective *sid-1(pk3321)* [*sid-1(-)*] mutants were used as a control for neuron-specific RNAi experiments. Two independent experiments were performed and pooled for analysis. See Table S4 for additional repeats and statistics.

**Figure S4. PMK-1 cascade signaling is required for enhanced pathogen resistance of *daf-2* mutants at young age**. (**A**-**B**) *pmk-1(km25)* [*pmk-1(-)*] mutations fully suppressed the enhanced PA14 resistance of young (day 1) adult *daf-2(e1370)* [*daf-2(-)*] animals (**A**), but partly suppressed the PA14 resistance of old (day 9) worms (**B**). (**C**-**D**) *nsy-1(ok593)* [*nsy-1(-)*] mutations fully suppressed the enhanced PA14 resistance of *daf-2(e1370)* [*daf-2(-)*] animals at day 1 adulthood (**C**), but partially suppressed that at day 9 adulthood (**D**). (**E**-**F**) *sek-1(km4)* [*sek-1(-)*] mutations decreased the PA14 resistance of wild-type (WT) and *daf-2(-)* worms at day 1 (**E**) and day 9 (**F**) adulthoods. See Table S5 for additional repeats and statistics.

**Figure S5. DAF-16/FOXO and HSF-1 are required for the enhanced pathogen resistance of *daf-2* mutants at old age**. (**A**-**B**) *daf-16(mu86)* [*daf-16(-)*] mutation fully suppressed the enhanced PA14 resistance of *daf-2(e1370)* [*daf-2(-)*] animals at day 1 (**A**) and day 9 (**B**) adulthoods. (**C**-**D**) *hsf-1(RNAi)* partly reduced the pathogen resistance of day 1 adult *daf-2(-)* animals (**C**), but fully suppressed that of day 9 adult *daf-2(-)* worms (**D**). (**E**-**F**) *skn-1* RNAi slightly decreased the PA14 resistance of wild-type (WT) and *daf-2(-)* worms at day 1 adulthood (**E**), but did not affect that at day 9 adulthood (**F**). See Table S5 for additional repeats and statistics.

**Figure S6. Several DAF-16/FOXO targets are further upregulated in aged *daf-2* mutants**. (**A**) Principal component analysis (PCA) of RNA seq data sets for wild-type (WT) and *daf-2(e1370)* [*daf-2(-)*] worms at day 1 and day 9 adulthoods (n = 3). (**B**) Representative PMK-1-regulated genes (T24B8.5, C17H12.5 and K08D8.5) were downregulated with age in WT and *daf-2(-)* worms. Error bars represent SEM (two-tailed Student’s *t*-test, n = 3) from RNA seq data. (**C**) mRNA levels of three selected DAF-16 targets, *mtl-1, hsp-16*.*2* and *sod-3*, in WT and *daf-2(-)* worms at day 1 and day 9 adulthoods in our RNA seq data (n = 3). (**D**-**E**) qRT–PCR analysis data confirmed that the mRNA levels of *mtl-1* (**D**) and *hsp-16*.*1/11* (**E**) were further upregulated with age in *daf-2(-)* compared to that in WT animals (n= 4). *ama-1* was used as a normalization control. (**F**-**G**) Venn diagram represents a significant overlap between genes whose expression levels were oppositely regulated during aging between WT and *daf-2(-)* animals (labeled as “DEGs with age”) and genes whose expression was negatively regulated by DAF-16/FOXO (Riedel et al., 2013) (**F**) and by HSF-1 (this study) (**G**) in *daf-2(-)* mutant backgrounds. RF: representation factor. *p* values were calculated with hypergeometric probability test.

**Figure S7. *ins-7* mutations enhanced immunocompetence at young and old ages**. *ins-7(tm2001)* [*ins-7(-)*^*2*^] mutation increased the survival of day 1 (**A**), day 6 (**B**), and day 9 (**C**) adult worms on PA14. See Table S5 for additional repeats and statistical analysis.

## Supplemental Table Legends

**Table S1. Genes whose expression was oppositely regulated with age between wild-type and *daf-2(-)* animals**. (**A**) Genes whose expression was age-dependently upregulated in wild-type but downregulated in *daf-2(-)* animals. (**B**) Genes whose expression was age-dependently downregulated in wild-type but upregulated in *daf-2(-)* animals.

**Table S2. Statistical analysis and additional repeats of immunosenescence assays with longevity mutants, or RNAi (related to Figure 1)**.

**Table S3. Statistical analysis and additional repeats of stress resistance assays with and without temporal *daf-2* RNAi treatment (related to Figures 2)**.

**Table S4. Statistical analysis and additional repeats of immunosenescence or lifespan assays using tissue-specific RNAi strains (related to Figure 2 and Supplemental Figures S2 and S3)**.

**Table S5. Statistical analysis and additional repeats of immunosenescence assays (related to Figures 3 through 7 and Supplemental Figures S4 through S7)**.

## Notes

### Competing Interest Statement

The authors have declared no competing interest.

