## Supplementary figures and images for "Reduction of insulin/IGF-1 receptor rejuvenates immunity via positive feedback circuit"

### Supplemental Figures

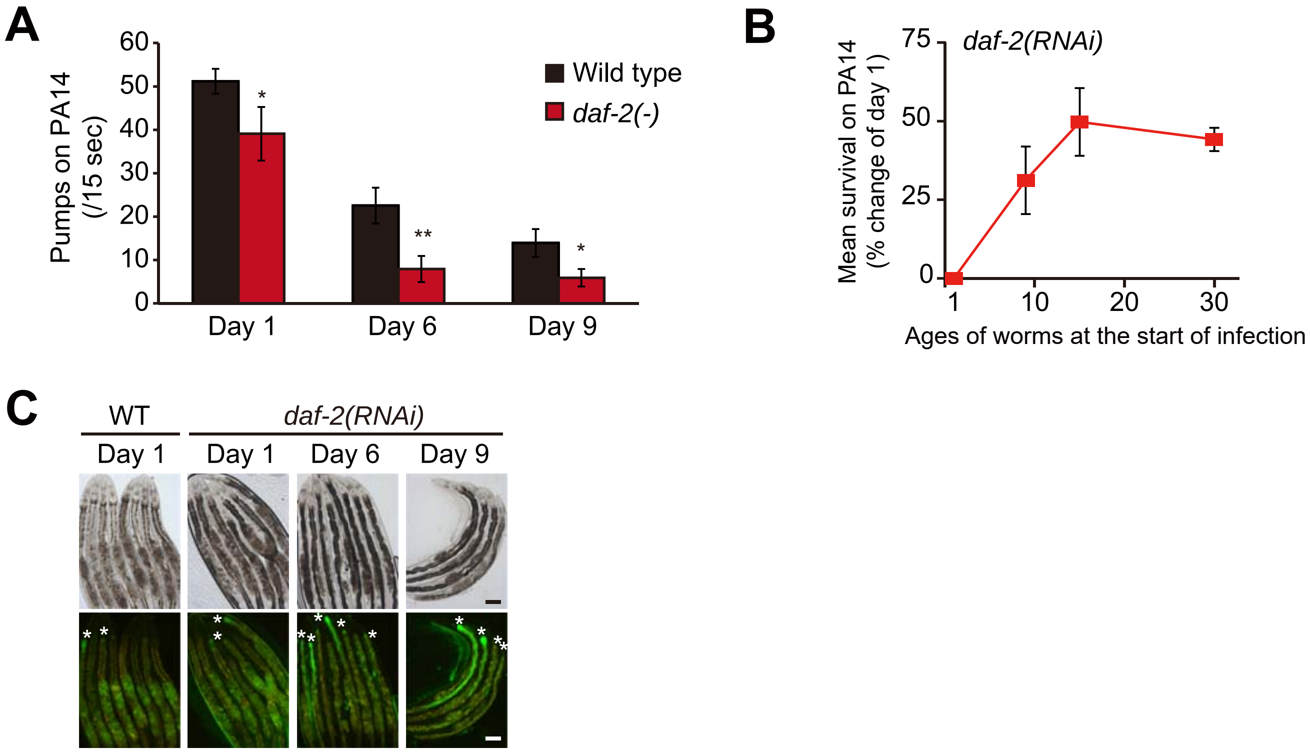

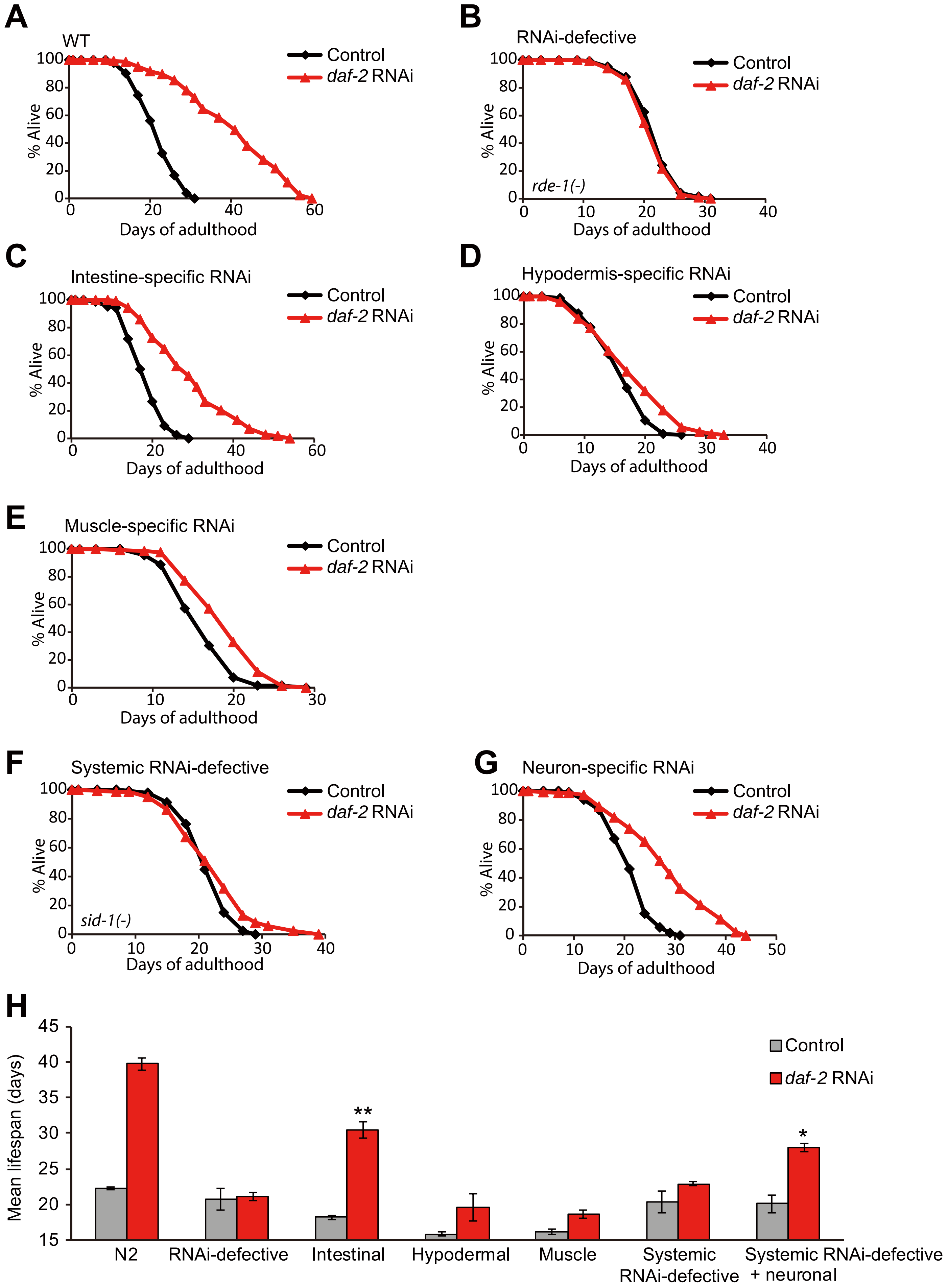

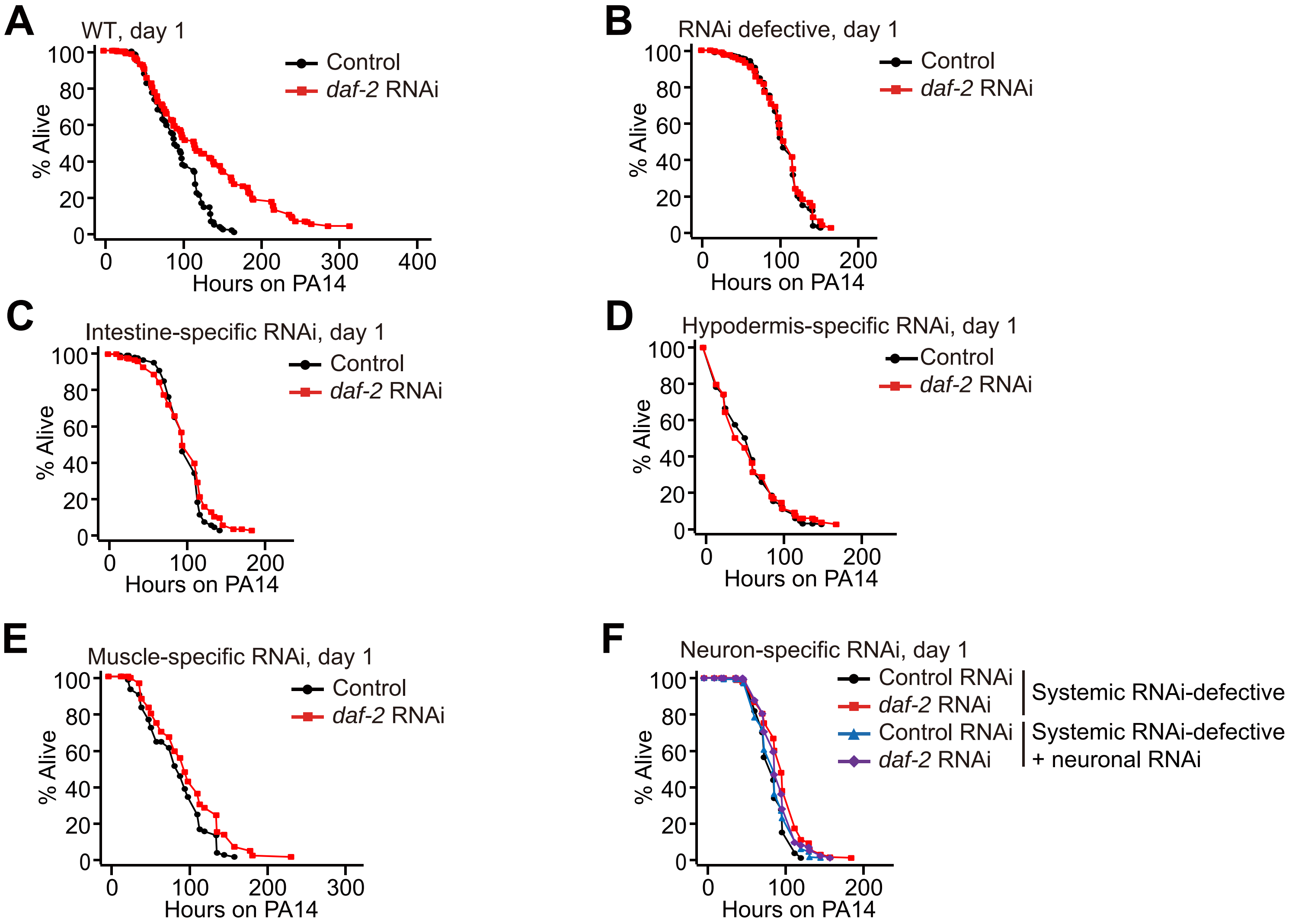

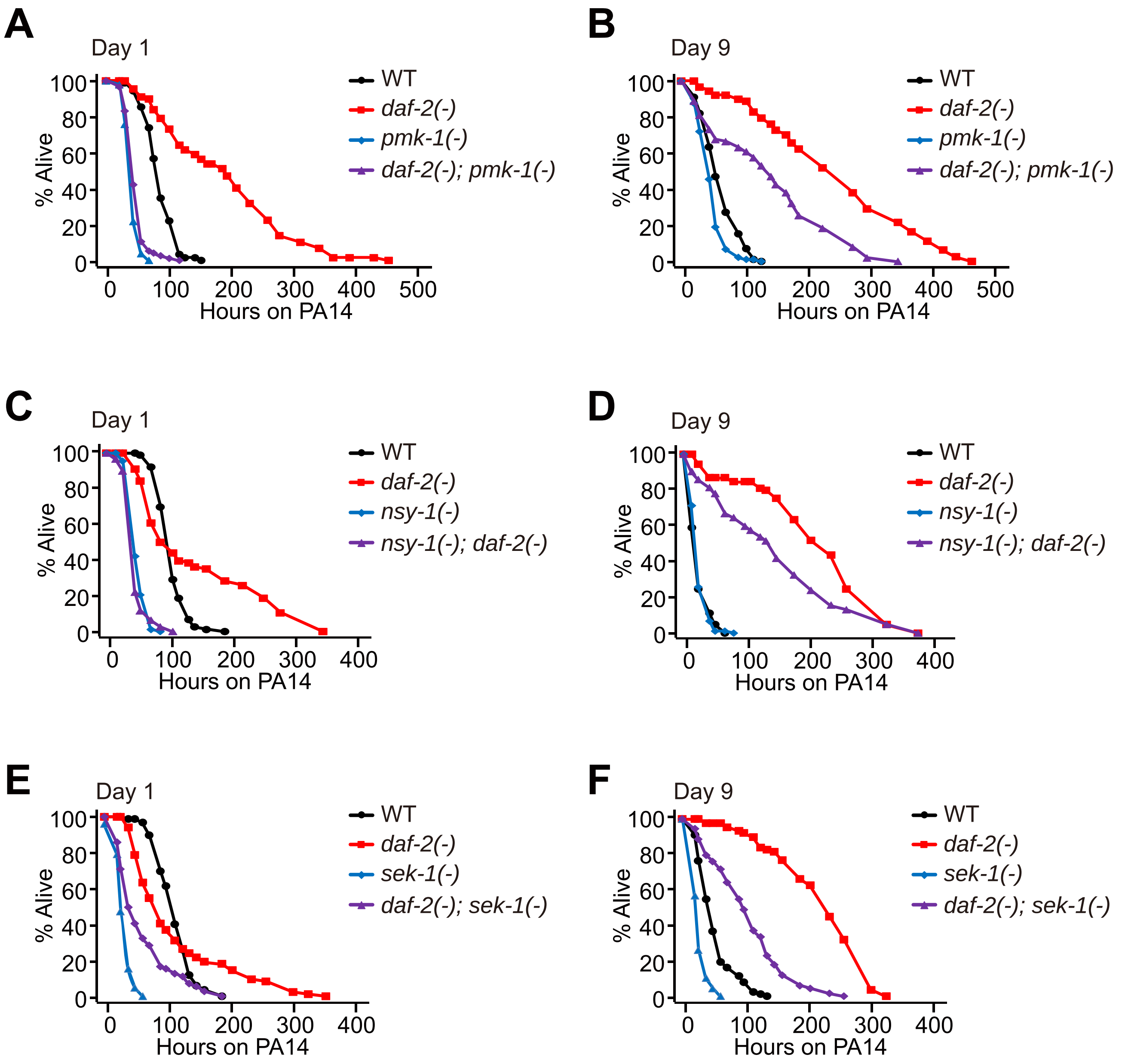

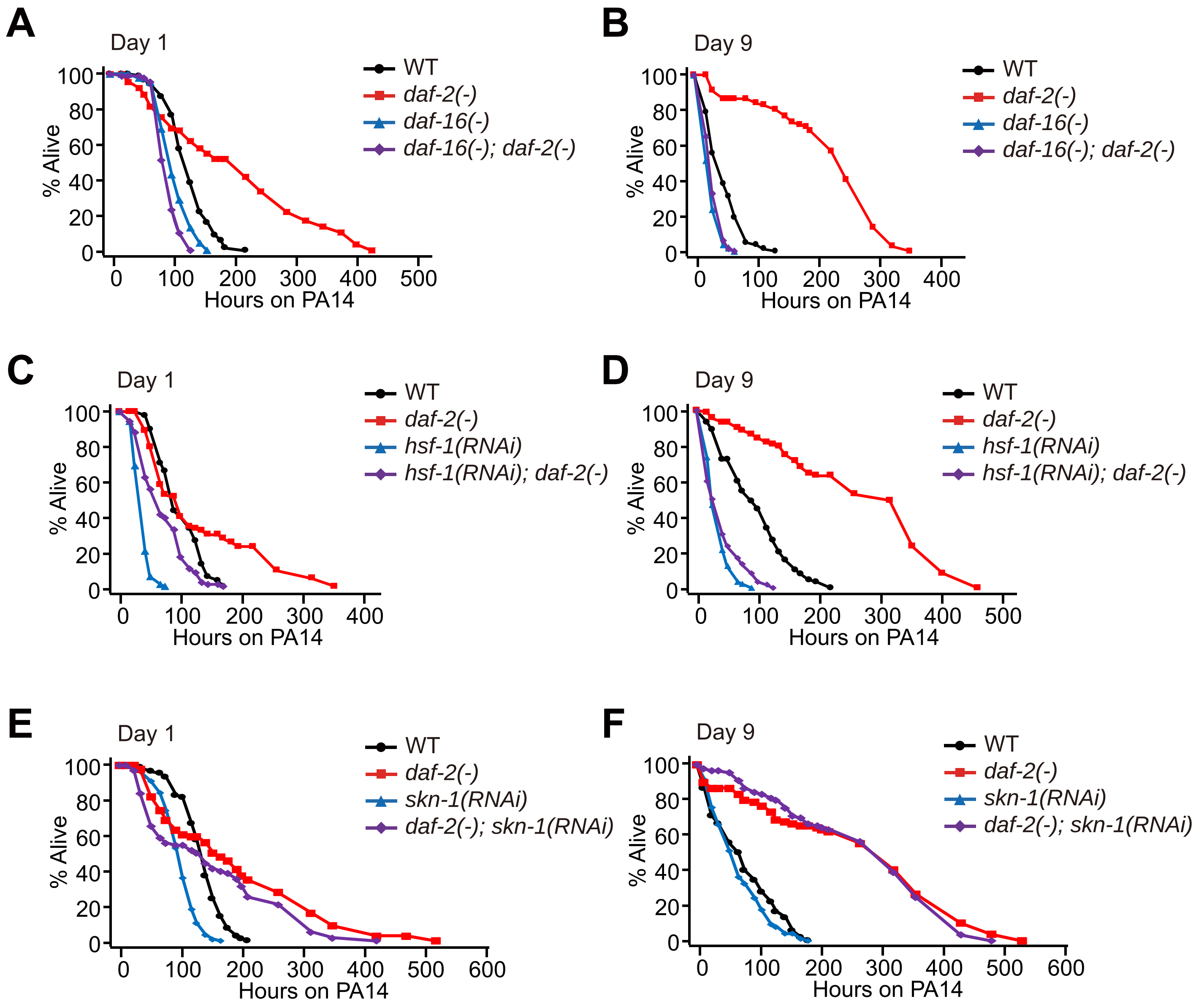

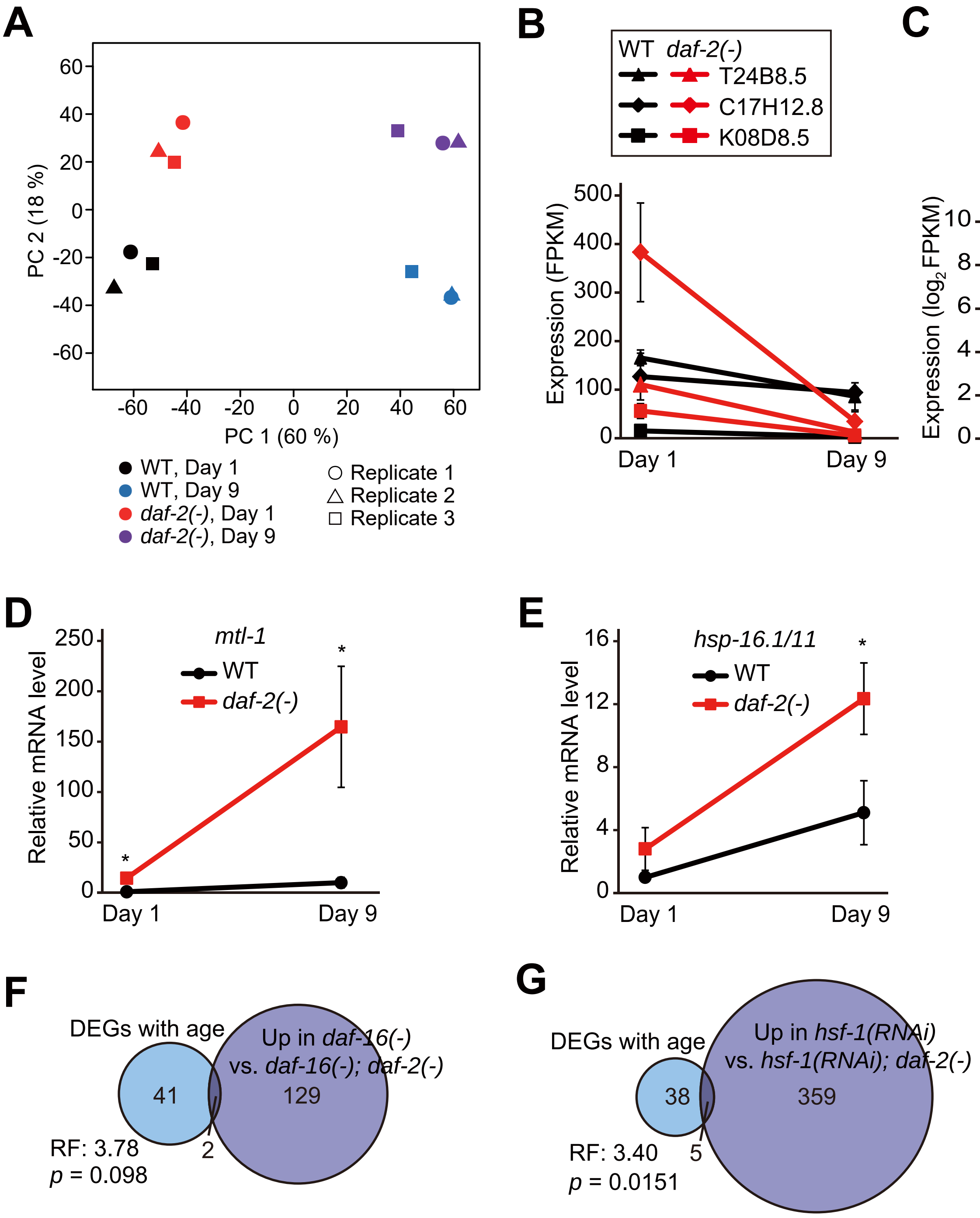

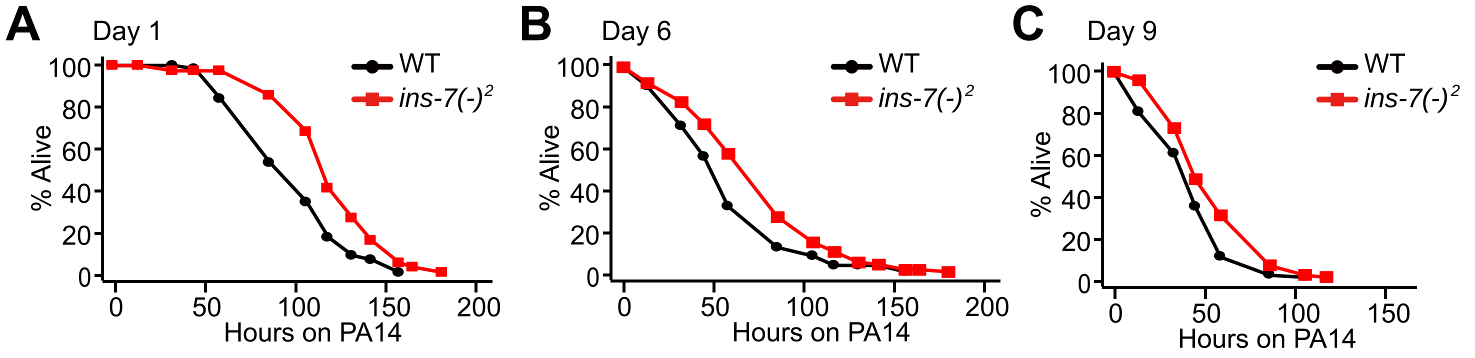
